# Neural stem cells traffic functional mitochondria via extracellular vesicles to correct mitochondrial dysfunction in target cells

**DOI:** 10.1101/2020.01.29.923441

**Authors:** Luca Peruzzotti-Jametti, Joshua D. Bernstock, Giulia Manferrari, Rebecca Rogall, Erika Fernandez-Vizarra, James C Williamson, Alice Braga, Aletta van den Bosch, Tommaso Leonardi, Ágnes Kittel, Cristiane Benincá, Nunzio Vicario, Sisareuth Tan, Carlos Bastos, Iacopo Bicci, Nunzio Iraci, Jayden A. Smith, Paul J Lehner, Edit Iren Buzas, Nuno Faria, Massimo Zeviani, Christian Frezza, Alain Brisson, Nicholas J Matheson, Carlo Viscomi, Stefano Pluchino

## Abstract

Neural stem cell (NSC) transplantation induces recovery in animal models of central nervous system (CNS) diseases. Although the replacement of lost endogenous cells was originally proposed as the primary healing mechanism of NSC grafts, it is now clear that transplanted NSCs operate via multiple mechanisms, including the horizontal exchange of therapeutic cargoes to host cells via extracellular vesicles (EVs).

EVs are membrane particles trafficking nucleic acids, proteins, metabolites and metabolic enzymes, lipids and entire organelles. However, the function and the contribution of these cargoes to the broad therapeutic effects of NSCs is yet to be fully understood. Mitochondrial dysfunction is an established feature of several inflammatory and degenerative CNS disorders, most of which are potentially treatable with exogenous stem cell therapeutics.

Herein we investigated the hypothesis that NSCs release and traffic functional mitochondria via EVs to restore mitochondrial function in target cells.

Untargeted proteomics revealed a significant enrichment of mitochondrial proteins spontaneously released by NSCs in EVs. Morphological and functional analyses confirmed the presence of ultrastructurally intact mitochondria within EVs (Mito-EVs) with conserved membrane potential and respiration. We found that the transfer of Mito-EVs to mtDNA-deficient L929 Rho^0^ cells rescued mitochondrial function and increased Rho^0^ cell survival. Furthermore, the incorporation of Mito-EVs into inflammatory professional phagocytes restored normal mitochondrial dynamics and cellular metabolism and reduced the expression of pro-inflammatory markers in target cells. When transplanted in an animal model of multiple sclerosis, exogenous NSCs actively transferred mitochondria to mononuclear phagocytes and induced a significant amelioration of clinical deficits.

Our data provide the first evidence that NSCs deliver functional mitochondria to target cells via Mito-EVs, paving the way for the development of novel (a)cellular approaches aimed at restoring mitochondrial dysfunction not only in multiple sclerosis, but also in degenerative neurological diseases.

## Introduction

Extracellular vesicles (EVs) are a heterogeneous population of secreted membrane vesicles with distinct biogenesis, biophysical properties and functions, which are virtually common to virtually all cells and life forms [27]. Despite their proven biological potential, the characterization and classification of this heterogeneous population of membrane vesicles has thus far been challenging.

A working basis for a consensus classification system of EVs is divided into three major subtypes based on biogenic, morphological, and biochemical properties: exosomes, microvesicles (MVs), and apoptotic bodies [64]. Exosomes are small vesicles, ranging 30 - 150 nm in diameter, generated from the inward budding of intracellular multi-vesicular bodies and released after the subsequent fusion with the plasma membrane [65]. MVs are membranous vesicles generated by clathrin-mediated shedding of the plasma membrane and released into the extracellular space, with a diameter ranging 50 - 1,000 nm [65]. Apoptotic bodies are generated through apoptotic fragmentation and blebbing with a resultant size range of 1,000 - 5,000 nm [18].

Neural stem cells (NSCs) are classically defined as a heterogeneous population of self-renewing, multipotent stem cells of the developing and adult central nervous system (CNS), which reside within specialized microenvironments and drive neurogenesis and gliogenesis [17, 44]. Data from our lab and peers have shown that in addition to the (expected) cell replacement, NSCs are strikingly able to engage in multiple mechanisms of action in the diseased CNS [3, 17], including the horizontal exchange of therapeutic cargoes to host cells via EVs [51]. However, the function and contribution of these cargoes to the broad therapeutic effects of NSCs is not fully understood.

We have recently focused on defining the nature and function of intercellular signalling mediated by EVs from NSCs [9, 26]. Using a series of computational analyses and high resolution imaging techniques, we have demonstrated that EVs deliver functional interferon gamma/interferon gamma receptor 1 (IFN-γ/Ifngr1) complexes to target cells [9]. We also discovered that EVs are endowed with intrinsic metabolic activities and harbour selective L-asparaginase activity catalysed by the enzyme Asparaginase-like protein 1 [26].

Recent evidence suggests that mitochondria play a key role in intercellular communications and that the release of mitochondria (or mitochondrial components) into the extracellular space has important functional consequences [63]. Growing attention has been given to the mechanisms regulating the mitochondrial exchange between cells. Several processes have been described, including the formation of actin-based tunnelling nanotubes (TNT) [31], cell-to-cell contact via gap junctions [28], and the release of EVs [45]. The latter mechanism seems to be central in regulating the exchange of mitochondria to inflammatory cells and could represent a novel mechanism of immunomodulation [4, 6, 30].

These pivotal observations prompted us to investigate whether NSC EVs also harbour organelles - including mitochondria - and what their functional relevance is for intercellular communication, immune modulation, and tissue repair.

Here, we used untargeted Tandem Mass Tag (TMT)-based proteomic analysis to investigate the protein content of EVs that are released by NSCs *in vitro*. We found an enrichment of mitochondrial proteins in both unfractionated EV preparations and exosome-specific fractions. Morphological and functional analyses unveiled structurally and functionally-intact, free and EV-embedded, mitochondria. We next studied the trafficking of these EV-embedded mitochondria (Mito-EVs) into target cells and found that Mito-EVs were efficiently incorporated by both somatic cells and professional phagocytes *in vitro*. Specifically, Mito-EVs rescue mitochondrial function in mtDNA deficient L929 Rho^0^ cells, as well as integrate into the host mitochondrial network of inflammatory macrophages (Mφ), thus modifying their metabolic profile and pro-inflammatory gene expression. When EVs and NSCs were used to treat mice with myelin oligodendrocyte glycoprotein (MOG)- induced experimental autoimmune encephalomyelitis (EAE), we found that exogenous NSCs actively transferred mitochondria to mononuclear phagocytes and induced a significant amelioration of clinical deficits.

Our data suggest that horizontal transfer of functional mitochondrial via EVs is a mechanism of signalling used by NSCs to modulate the physiology and metabolism of target cells, opening a possible new avenue for the development of acellular therapies aimed at correcting mitochondrial dysfunction in the CNS.

## Materials and Methods

### Neural Stem Cells (NSCs)

Somatic NSCs were obtained from the subventricular zone (SVZ) of 7-12-week-old (18-20 g) C57BL/6 mice (Charles River, UK). Briefly, mice were humanely culled by cervical dislocation followed by decapitation, the parietal bones were cut cranially to caudally using micro-chirurgic scissors, and the brains removed. A brain slice matrix was used to obtain 3 mm thick brain coronal sections starting from 2 mm after the anterior pole of the brain. The SVZ of the lateral ventricles was isolated from coronal sections using iridectomy scissors. Tissues derived from at least 2 mice were pooled. Dissected tissues were transferred to a 15 ml tube with digestion medium [early balance salt solution (EBSS, Gibco), papain (1 mg/ml, Worthington), ethylenediaminetetraacetic acid (EDTA) (0.2 mg/ml, Sigma-Aldrich) and L-cysteine (0.2 mg/ml, Sigma-Aldrich)] and incubated for 45 min at 37°C on a rocking platform. At the end of the incubation, the tube was centrifuged at 200 x *g* for 12 min, the supernatant was removed and the pellet was mechanically disaggregated with 2 ml of EBSS. The pellet was centrifuged again at 200 x *g* for 12 min and then dissociated with a 200 µl pipette and seeded in complete growth medium (CGM). CGM was constituted of mouse NeuroCult basal medium (Stem Cell Technologies) plus mouse NeuroCult proliferation supplements (Stem Cell Technologies) added with 2 µg/ml heparin (Sigma-Aldrich), 20 ng/ml epidermal growth factor (EGF) and 10 ng/ml basic fibroblast growth factor (bFGF). After approximately 4-7 days, a small percentage of the isolated cells begun to proliferate, giving rise to small cellular aggregates (i.e. neurospheres). When neurospheres reached the necessary dimension (150-200 µm diameter), the cells were harvested in a 15 ml tube and centrifuged at 100 x *g* for 8 min. The supernatant was then removed, and the pellet dissociated by enzymatic digestion with Accumax at 37°C for 10 min. The number of viable cells was determined by trypan blue exclusion and viable cells were re-seeded at clonal density 8,000 cells/cm^2^. Mycoplasma tested (negative) NSCs at passage n ≤ 30 were used in all experiments.

### Extracellular Vesicle (EV) and Exosome isolation

For EV isolation, NSCs were single-cell dissociated and plated at a concentration of 12×10^6^ cells/10 ml medium/T75 culture flask. After 18 hrs, EVs were isolated as previously described [9]. Briefly, supernatants were collected and centrifuged for 15 min at 300 x *g* to remove cellular pellets. The supernatant was collected and centrifuged for 15 min at 1,000 x *g* to remove cellular debris. The supernatant was then collected and subjected to ultracentrifugation at 100,000 x *g* for 70 min at 4°C using an Optima XPN-80 ultracentrifuge with SW 32 Ti swinging rotor (Beckman Coulter). Pellets were washed in PBS 1X and EVs were subjected to an additional centrifugation at 100,000 x *g* for 30 min at 4°C using an Optima MAX ultracentrifuge, with TLA110 fixed angle rotor (Beckman Coulter).

For exosome isolation, pellets of EVs were resuspended in 0.5 ml 0.32 M sucrose. The solution was layered on a 10 ml continuous sucrose density gradient [0.32–2 M sucrose, 5 mM HEPES (pH 7.4)] and centrifuged for 18 hrs at 100,000 x *g* (SW 32.1 Ti; Beckman Coulter) with no brake. Fractions were collected from the top (low density) to the bottom of the tube (high density) of the sucrose gradient. Single fractions from the gradient were harvested, diluted in PBS 1x, centrifuged at 100,000 g for 70 min at 4°C using an Optima XPN-80 ultracentrifuge with SW 32 Ti swinging rotor (Beckman Coulter) and then processed for further analyses.

### Alternative EV isolation protocols

EV isolation with Qiagen (cat. No. 76743) and Invitrogen (cat. No. 4478359) commercially available kits was carried out following manufacturers’ instructions. EVs isolation using the method of Wang et al. 2017 was performed as previously described [68]. Briefly, supernatants from cultured NSCs (as described above) were collected, transferred to 50 ml polypropylene centrifuge tubes and centrifuged for 10 min at 300 x *g*. The supernatants were then collected and transferred into new 50 ml polypropylene centrifuge tubes before being subject to further centrifugation for 30 min at 2000 x *g*. Subsequently supernatants were transferred to 100 ml polycarbonate tubes and centrifuged for 20 min at 16,500 x *g* Supernatants were then filtered through a 500 ml disposable filter unit (0.22 μm) and transferred to new 100 ml polycarbonate tubes prior to centrifugation for 70 min at 120,000 x *g*. Pellets resulting from this step are composed by EVs. Pellets were let to dry and immediately resuspend in 1ml of ice-col PBS; pellets were then pooled together in ultracentrifuge tubes and centrifuged for 70 min at 100,000 x *g*. Supernatants were completely removed and pellets were processed for protein analysis.

### Mitochondria-enriched preparation (Mito)

Mitochondria were isolated either from NSCs plated at 12×10^6^ cells in 10 ml (high density) or from NSC plated at 1.5×10^6^ cells in 10 ml (standard density), as previously described[15]. Briefly, cells were collected and centrifuged at 600 x *g* for 8 min. NSCs pellet was washed two times with cold PBS and resuspended in one cell pellet volume of ice cold 0.1X homogenisation medium IB (IB10X: 0.35M Tris–HCl, pH 7.8, 0.25 M NaCl, and 50 mM MgCl_2_). Cells were then homogenized at 1,600 rpm for 3 min with a Teflon pestle and 10X IB was immediately added at 1/10 of the initial cells pellet volume, to make the medium isotonic. The homogenate was then centrifuged at 1600 x *g* for 3 min at 4°C to pellet unbroken cells, debris and nuclei. Supernatant was then collected and centrifuged at 13,000 rpm for 1 min to pellet down mitochondrial enriched pellet. Once isolated, mitochondrial pellet was washed with homogenisation medium 1X IB and centrifuged 1,300 rpm, 1 min. Mitochondrial pellet was then resuspended in 200 µl of Medium A (0.32M sucrose, 1mM EDTA, and 10 mM Tris–HCl), centrifuged down and resuspended in the appropriate volume of medium/PBS for downstream utilization.

### Nanoparticle tracking analysis (NTA) and tunable resistive pulse sensing (TRPS) analysis

EVs derived by 36×10^6^ of NSCs were diluted respectively 1:1000 and 1:500 with PBS 1X for NTA analysis using a Nanosight NS500 (Malvern Instruments Ltd) fitted with an Electron Multiplication Charge-Couple Device camera and a 532 nm laser. At least 3 videos were recorded for each sample using static mode (no flow). Between each capture the sample was advanced manually and the temperature was monitored and maintained at 25±1°C. Data analysis was carried out on NTA 3.2 software using a detection threshold between 5 or 6.

The concentration and size distribution of EVs was also analysed with TRPS (qNANO, Izon Science Ltd) using a NP150 Nanopore at 0.5 V with 47 mm stretch. The concentration of particles was standardized using 100 nm calibration beads (CPC 100) at a concentration of 1×10^10^ particles/ml.

### Proteomic analysis

Sample preparation: EVs and exosomes were purified from murine NSCs culture supernatants by ultracentrifugation (total particles) followed by density gradient centrifugation (exosomes) as previously described[9]. For comparison with NSCs whole cell lysates (NSCs), washed NSCs cell pellets were processed in parallel. All samples were prepared in triplicate (biological replicates). Tryptic digests were made using an IST-NHS sample preparation kit (Preomics GmBH) according to the manufacturer’s instructions with minor modifications. Briefly, samples were solubilised in proprietary lysis buffer, and sonicated 10 times 30s on/off in a Bioruptor sonicator (Diagenode). Lysates were diluted 10-fold and quantified by BCA assay against a BSA standard curve in diluted lysis buffer. Digestion was performed at 37°C for 3hrs. Tandem Mass Tag (TMT) labelling was performed on the digestion columns as per the manufacturer’s instructions. After elution, TMT labelling of at least 98% peptides was confirmed for each sample before pooling and subjecting to high pH reversed-phase (HpRP) fractionation. This was conducted on an Ultimate 3000 UHPLC system (Thermo Scientific) equipped with a 2.1 mm × 15 cm, 1.7µ Acquity BEH C18 column (Waters, UK). Solvent A was 3% acetonitrile (ACN), solvent B was 100% ACN, solvent C was 200 mM ammonium formate (pH 10). Throughout the analysis solvent C was kept at a constant 10%. The flow rate was 400 µl/min and UV was monitored at 280 nm. Samples were loaded in 90% A for 10 min before a gradient elution of 0–10% B over 10 min (curve 3), 10-34% B over 21 min (curve 5), 34-50% B over 5 min (curve 5) followed by a 10 min wash with 90% B. 15 s (100 µl) fractions were collected from the start of the gradient elution. Fractions were pooled orthogonally to generate a final 24 fractions.

Mass spectrometry: All samples were resuspended in 5% dimethyl sulfoxide/0.5% trifluoroacetic acid. Sample was analysed using a nanoLC-MS platform consisting of an Ultimate 3000 RSLC nano UHPLC (Thermo/Fisher Scientific) coupled to an Orbitrap Fusion (Thermo/Fisher Scientific) instrument. Samples were loaded at 10 μl/min for 5 min onto an Acclaim PepMap C18 cartridge trap column (300 µm x 5 mm, 5 µm particle size) in 0.1% TFA. After loading a linear gradient of 3-32% solvent B over 180 min was used for sample separation over a column of the same stationary phase (75 µm x 75 cm, 2 µm particle size) before washing at 95% B and equilibration. Solvents were A: 0.1% formic acid (FA) and B:100% ACN/0.1% FA. Electrospray ionisation was achieved by applying 2.1kV directly to a stainless-steel emitter tip. Instrument settings were as follows. MS1: Quadrupole isolation, 120’000 Resolution, 5e5 AGC target, 50 ms maximum injection time, ions injected for all parallisable time. MS2: Quadrupole isolation at an isolation width of m/z 0.7, CID fragmentation (NCE 30) with ion trap scanning out in rapid mode from m/z 120, 5e3 AGC target, 70ms maximum injection time, ions accumulated for all parallisable time in centroid mode. MS3: In Synchronous precursor selection mode the top 10 MS2 ions were selected for HCD fragmentation (65NCE) and scanned out in the orbitrap at 50’000 resolution with an AGC target of 2e4 and a maximum accumulation time of 120 ms, ions were not accumulated for all parallelisable time. For all experiments the entire MSn cycle had a target time of 3s.

Data processing and analysis: Spectra were searched by Mascot within Proteome Discoverer 2.2 in two rounds of searching. The first search was against the UniProt Mouse reference proteome and a compendium of common contaminants (GPM). The second search took all unmatched spectra from the first search and searched against the Mouse trEMBL database. Search parameters were as follows. MS1 Tol: 10 ppm, MS2 Tol: 0.6 Da, Fixed mods: Carbamidomethyl (C) and TMT (N-term, K), Var mods: Oxidation (M), Enzyme: Trypsin (/P). For HCD-OT Experiments. MS1 Tol: 10 ppm, MS2 Tol: 0.05 Da, Fixed mods: Carbamidomethyl (C) and TMT (N-term, K), Var mods: Oxidation (M), Enzyme: Trypsin (/P). MS3 spectra were used for reporter ion-based quantitation with a most confident centroid tolerance of 20 ppm. PSM FDR was calculated using Mascot percolator and was controlled at 0.01% for ‘high’ confidence PSMs and 0.05% for ‘medium’ confidence PSMs. Normalisation was automated and based on total s/n in each channel. Proteins/peptides satisfying at least a ‘medium’ FDR confidence were taken forth for further analysis. To compare protein abundances in particles, exosomes and NSCs, moderated *t*-tests were performed using the *limma* R/Bioconductor software package, with false discovery rate (FDR)-adjusted p values (q values) calculated according to the Benjamini-Hochberg method. To analyse subcellular localisations of proteins identified or enriched in particles and/or exosomes, Gene Ontology Cellular Component (GOCC) terms were imported using the Perseus software platform. Further data manipulation and general statistical analysis were conducted using Excel and XLSTAT. The proteomic data described in this study have been deposited to the ProteomeXchange consortium via the PRIDE partner repository (accessible at http://proteomecentral.proteomexchange.org). A dataset identifier will be provided prior to publication, and an access key made available to reviewers on request.

### Western blotting

Whole cell pellets were solubilised in 100 μl of RIPA buffer (10 mM Tris HCl pH 7.2, 1% v/v sodium deoxycholate, 1% v/v Triton X-100, 0.1% v/v sodium dodecyl sulfate (SDS), 150 mM NaCl, 1 mM EDTA pH 8) in presence of Complete Protease Inhibitor Cocktail (Roche) and Halt Phosphatase Inhibitor Cocktail (Pierce). EVs, exosomes and isolated mitochondria pellets were solubilised in 50 μl of RIPA buffer with 3% v/v SDS. Protein were then quantified using Bio-Rad DC Protein Assay Kit II.

Protein extract were then loaded onto NuPAGE LDS sample buffer 4X (Invitrogen) under reducing (sample reducing agent 10X) and non-reducing conditions and separated by SDS-PAGE on 4-12% precast NuPAGE Bis/Tris gels (1.5 mm thickness). Before loading, samples were heated at 95°C for 5 min, then loaded onto the gels and then run at 120V in MOPS running buffer. Samples were then transferred on polyvinylidene fluoride membranes (0.45 μm pore size, Immobilon) filter paper sandwich using XCell II Blot Module and NuPAGE transfer buffer (Invitrogen).

For immunoblot analysis, the membranes were blocked with 5% non-fat milk in 0.1% PBS-Tween 20 (Sigma) for 1 hr at room temperature and then incubated with the following primary antibodies (diluted in 5% non-fat milk in 0.1% PBS-Tween 20) for 18 hrs at 4°C: Mouse monoclonal Total OXPHOS rodent WB cocktail (Abcam ab110413, 1:1,000); mouse monoclonal Total OXPHOS blue-native WB cocktail (Abcam ab110412, 1:1,000); rat monoclonal anti-CD9 (BD Pharmingen 553758, 1:1000); mouse monoclonal anti-Pdcd6ip (AIP-1/Alix) (BD transduction lab 611620, 1:500); goat polyclonal anti-TSG101 (Santa Cruz sc-6037, 1:500); rabbit monoclonal TOMM20 (Santa Cruz sc-11415,1:1000); rabbit polyclonal ant-H3 (Abcam ab1791,1:10,000), mouse monoclonal anti-ß-actin (Sigma, A1978 1:10000), mouse monocolonal anti-Gm130/Golga2 (BD Transduction Laboratories 610823, 1:1,000). Molecular weight marker: SeeBlue Plus2 (Invitrogen).

After primary antibody incubation, membranes were washed 3 times for 10 minutes with 0.1% PBS-Tween 20 and incubated with the appropriate horseradish-peroxidase-conjugated secondary antibodies (Thermo Scientific) for 1 hr at room temperature. Amid each primary antibody incubation, membranes were subjected to stripping protocol. Briefly, after image acquisition, the membrane was washed with PBS 1x and then incubated for 1hr with stripping buffer (10x: 75.08 g Glycine 410225 Sigma, 10 g SDS 862010 Sigma, 1L dH_2_O, pH 7.4). After 2 washes with 0.1% PBS-Tween 20 the membrane was then blocked with 5% non-fat milk in 0.1% PBS-Tween 20 plus for 1 hr and then incubated with the next relevant primary antibodies (diluted in 5% non-fat milk in 0.1% PBS-Tween 20). The secondary antibodies used for all the WBs were: goat anti rabbit-HRP (31460, Thermo Scientific, 1:10,000), goat anti mouse-HRP (31430, Thermo Scientific, 1:20,000), rabbit anti-goat (31402, Thermo Scientific, 1:10,000), goat anti-rat (31470, Thermo Scientific, 1:10,000).

Immunoreactivity was revealed by using Western Lightning® Plus-ECL Prime Western Blotting Detection Reagent (Amersham, RPN2232) according to manufacturer’s instruction. Image were acquired on a Biorad Chemidoc MP system, using high sensitivity acquisition in signal accumulation mode.

### Genomic PCR

Total DNA was extracted from cells pellet derived from 3×10^6^ NSCs and from L929 Rho^0^, EVs derived from 12 ×10^6^ NSCs, isolated mitochondria derived from 12×10^6^ NSCs. Prior to DNA isolation, samples were resuspended in PBS suspensions and pre-treated with DNase I (Sigma, #D5025) [50 U/ml] for 1 hr at 37°C to remove possible external contaminating DNA and RNA. Nuclease reaction was stopped by PBS wash, at the appropriate centrifugation speed for each sample (isolated mitochondria preparation: 12,400 x *g* for 1 min; NSCs: 16,000 x *g* for 5 min; EVs: 100,000 x *g* for 45 min, the ultracentrifugation steps was run with TLA110 rotor-Beckman Instruments). DNase I pre-treated samples were all resuspended in 200 μl of PBS and total DNA was finally extracted with DNeasy Blood & Tissue Kit (Qiagen) according to manufacturer’s instructions. Total DNA yield and purity of the extracted DNA were determined by using a Nanodrop spectrophotometer (Thermo Scientific, Waltham, MA).

Nuclear and mitochondrial DNA (nDNA and mtDNA) content of each sample was assessed by polymerase chain reaction (PCR) amplification of the mtDNA gene *NADH dehydrogenase subunit 1* (*mt-ND1*) and the nuclear DNA gene *succinate dehydrogenase complex subunit D* (*Sdhd*), starting from 25 ng of total DNA extract and using KAPA biosystem 2G kit according to manufacturer’s instructions. Primer sequences: *mt-ND1* - F: 5’-TGCACCTACCCTATCACTCA-3’; R: GGCTCATCCTGATCATAGAATGG; expected product: 148 bp; *Sdhd* - F: CTTGAATCCCTGCTCTGTGG; R: AAAGCTGAGAGTGCCAAGAG; expected product: 1660 bp. Amplification reaction parameters: initial denaturation at 95°C for 3 min (1x); denaturation 95°C for 15 sec, annealing temperature 60°C for 15 sec, elongation time ET 72°C for 1 sec (all 35x); final extension 72°C for 1 min. Amplified nucleic acids (25 μl) were mixed with Purple dye loading buffer 6X (Biolegends) loaded onto a 2% Agorose/Gel Red (Biolegends)/1x TAE gel and run for 40 min at 120V. Loading Buffer: TAE 1X electrophoresis buffer. At run completion, DNA bands were visualized with UV-light (312 nm) with Biorad Chemidoc MΦ system, and directly photographed.

### Transmission Electron Microscopy (TEM) and Cryo-Transmission Electron Microscopy (Cryo-TEM)

For TEM analysis EVs were derived from NSCs, as described above. Supernatant was carefully removed and the pellets were fixed with PFA 4%-PBS1X for 10 min at room temperature, and then in fresh portion of PFA 4%-PBS1X for overnight at 4°C. After removal of the fixative, and a short rinse with PBS1X, the pellets were post-fixed in 1% OsO_4_ (Taab) for 30 min, rinsed with distilled water, dehydrated in graded ethanol, including block staining with 1% uranyl-acetate in 50% ethanol for 30 min, and embedded in Taab 812 (Taab). Overnight polymerization at 60 °C was followed by sectioning, and the ultrathin sections were analysed using a Hitachi 7100 electron microscope (Hitachi Ltd) equipped with Veleta, a 2000 × 2000 MegaPixel side mounted TEM CCD camera (Olympus).

For immuno-gold labelling and cryo-transmission electron microscopy (cryo-TEM) analysis, 10-nm gold nanoparticles (NP) were conjugated with anti-TOMM20 monoclonal antibody (Abcam ab232589) following the procedures previously described[2]. Re-suspended EV pellets were labelled for 1 h with anti-TOMM20-gold-NP at 1-3 × 10^15^ gold-NP/L. Immuno-gold labelled EV samples were processed for cryo-TEM according to standard procedures. A 4 μL aliquot was deposited on an EM grid coated with a perforated carbon film. After draining the excess liquid with a filter paper, grids were plunge-frozen into liquid ethane cooled by liquid nitrogen using a Leica EMCPC cryo-chamber. For cryo-TEM observation, grids were mounted onto a Gatan 626 cryoholder and transferred to a Tecnai F20 microscope (ThermoFisher, USA) operated at 200 kV. Images were recorded with an Eagle 2k CCD camera (FEI, USA).

### Blue Native Polyacrylamide Gel Electrophoresis (BN-PAGE)

Extractions of proteins in native conditions from EVs, isolated mitochondria and NSCs were performed as previously described [47].

EVs native protein extraction: EVs derived from 36×10^6^ NSCs were incubated on ice for 10 min in hypotonic buffer solution (EDTA-Na 1 mM, Tris HCl 6 mM, pH 8) with added Protease Inhibitor Cocktail 1X. Next, EVs were homogenized with a pestle and isotonic conditions were restored by adding sorbitol 1 M (final concentration 0.32 M), EDTA 0.5 M (final concentration 10 mM) and Tris-HCl 1 M (final concentration 10 mM). The extract was then centrifuged at 10,000 x *g* for 10 min, washed with PBS 1X and centrifuged again at 10,000 x *g* for 5 min. Pellet was resuspended with Medium A supplemented with bovine serum albumin (BSA) and centrifuged 10,000 x *g* for 10 min. This pellet was then resuspended in 50 μl of Medium A and 5 µl were taken for BCA protein quantification. Ultimately, the extracts were centrifuged again at 10,000 x *g* for 10 min and resuspended at a protein concentration of 5 mg/ml in solubilisation buffer (1.5 M aminocaproic acid, 50 mM Bis-Tris/HCl pH= 7.0). For the solubilisation, 1.6 mg of n-dodecyl-β-D-maltoside (DDM) per mg of protein was added. Samples were incubated on ice for 5 minutes, then centrifuged at 20,000 x *g* for 30 min at 4°C. Supernatants were collected and a same volume of sample buffer (750 mM aminocaproic acid, 50 mM Bis–Tris/HCl, pH 7.0, 0.5 mM EDTA, and 5% Serva Blue G) was added. The samples were stored at −80°C until it was time to perform BN-PAGE.

NSCs native proteins extractions: pellet from 3×10^6^ cells was resuspended in 200 μl of PBS 1X. 200 μl of cold digitonin solution (8 mg/mL dissolved in PBS), was added to the sample and kept in ice. After 10 min of incubation, digitonin was diluted by adding 1 ml of cold PBS and cells were centrifuged at 10,000 x *g* for 5 minutes at 4°C. This step was repeated twice. Cells pellets were then resuspended in 100 μl of PBS and 5 μl was taken for protein quantification. Next, PBS was removed by centrifuging at 10,000 x *g* for 5 min and cell pellet was solubilised as described for EV samples.

Isolated mitochondria native protein extraction: mitochondrial pellet was resuspended in 100 μl of PBS and 5 μl was taken for protein quantification. Next, Medium A was removed by centrifuging at 10,000 x g for 5 min, and the pellet containing the isolated mitochondria, was solubilized and processed as previously described for EVs and NSCs.

A total of 50 μg of protein for each of the samples was loaded into a pre-cast NativePAGE 3-12% Bis-Tris gel (Invitrogen). For the run, 1x NativePAGE Running Buffer plus 1x NativePAGE Cathode Buffer Additive (Invitrogen) was added to the cathode and 1x NativePAGE Running Buffer was added to the anode. Half-way through the run the cathode buffer was substituted for 1x NativePAGE Running Buffer plus 0.1x NativePAGE Cathode Buffer Additive and run until the front reached the end of the gel.

### In-gel mitochondrial complexes activity

After the electrophoresis gels were incubated with the following buffers containing the substrates and electron acceptors necessary for the colorimetric reactions to take place [70].

Complex I: 1 mg/ml nitro blue tetrazolium (NBT), 1 mg/ml NADH in 5 mM Tris-HCl, pH 7.4.

Complex II: 1 mg/ml nitro blue tetrazolium (NBT), 20 mM sodium succinate, 0.2 mM phenazine methasulfate (PMS) in 5 mM Tris-HCl, pH 7.4

Complex IV: 1 mg/ml 3,3’-diaminobenzidine tetrahydrochloride (DAB), 24 U/ml catalase, 1 mg/ml cytochrome c, 75 mg/ml sucrose in 50 mM potassium phosphate buffer, pH 7.4.

### Mitochondrial Membrane potential measurement

Mitochondrial membrane polarization was determined using MitoProbeTM JC-1 Assay Kit for Flow Cytometry (M34152 ThermoFisher) according to manufacturer’s instructions. For JC-1 (5,5,6,6′-tetrachloro-1,1′,3,3′-tetraethylbenzimidazolylcarbocyanine iodide) staining 6×10^6^ NSCs, as well as EVs and mitochondria derived from 6×10^6^ NSCs were resuspended in 500 μl of NSCs medium and loaded with JC-1 (2 μM), and incubated for 30 min at 37°C, CO_2_ 5%. After incubation samples were washed once, resuspended in 400 µl 1x PBS, plated in quadruplicate (100 μl/1M/well) in a 96-well flat bottom plate and immediately analysed on a Tecan Infinite M200 Pro plate reader. J-aggregates red fluorescence was read at Ex488/Em590, while monomers green fluorescence at Ex488/Em530. Blank signal from PBS1X only were subtracted from samples fluorescence values and finally data were expressed as JC1 ratio Red-over-Green fluorescence (JC1-R/G).

### High resolution respirometry (HRR - Oroboros oxygraph respiration assay)

Mitochondrial respiration in permeabilized NSCs and hypotonic-shock treated EVs was analysed by high resolution respirometry (HRR) (Oroboros oxygraph, Innsbruck, Austria). HRR analysis was performed based on previously published methods[22, 42]. Briefly, a 2 ml oxygraph (Oxygraph-2k; Oroboros) chamber was washed with 70% ethanol, rinsed 3 times with distilled water, then filled with the respiration medium (Medium A plus 1 mM adenosine diphosphate (ADP), 2 mM potassium phosphate and 1 mg/ml fatty acid free BSA) to be used in each of the assays. 20×10^6^ NSCs were resuspended in 2 ml of Medium A (20 mM HEPES (adjusted to pH 7.1 with NaOH or KOH), 250 mM sucrose, 10 mM MgCl_2_). For permeabilization, the cell suspension was incubated with 5 μl of 1% digitonin solution for 1 min at room temperature on a tube oscillator. Digitonin was then diluted with 5 mL of Medium A and removed by centrifugation at 1000xg for 3 min. Finally, the pellet was resuspended in 2.1 ml of respiration medium and added into the chamber.

EVs from 36 ×10^6^ were isolated and subjected to hypo-osmotic shock as described above. Upon removal of homogenization Medium A through centrifugation at 10,000 x *g*, for 10 min, hypotonic-shock treated vesicles were equilibrated in MAITE medium (25 mM sucrose, 75 mM sorbitol, 100 mM KCl, 0.05 mM EDTA, 5 mM MgCl_2_, 10 mM Tris-HCl, 10 mM phosphate, pH 7.4). The suspension was centrifuged at 10,000 x *g* for 10 min, resuspended in 100 μl of MAITE buffer, then diluted up to 2.1 mL of MAITE plus 1 mg/ml fatty acid free BSA.

Once samples were loaded in the chambers, a polyvinylidene fluoride stopper was inserted to generate a closed system with a final volume of 2 ml. Oxygen concentration was recorded at 0.5 Hz and converted from voltage to oxygen concentration using a two-point calibration. Respiration rates (O_2_ flux) was calculated as the negative time derivative of oxygen concentration (Datlab Version 4.2.1.50, Oroboros Instruments). The O_2_ flux values were corrected for the small amount of back-diffusion of oxygen from materials within the chamber, any leak of oxygen from outside of the vessel, and oxygen consumed by the polarographic electrode. A protocol involving serial additions of selected mitochondrial complex substrates and inhibitors was performed for a comprehensive assessment of mitochondrial function. The following substrates and inhibitors were progressively injected with Hamilton syringe at the respective concentrations: 5 mM glutamate and 5 mM malate (NADH-linked substrates); 0.1 μM rotenone (complex I inhibitor); 5 mM glycerol-3-phosphate and 5 mM succinate (FADH2-linked substrate); cytochrome *c* (Cyt c) to compensate for a possible loss due to outer membrane disruption; 20 nM antimycin A (complex III inhibitor); 1 mM *N, N, N′, N′*- tetramethyl-*p*-phenylenediamine (TMPD, complex IV electron donor) and finally 0.1 mM KCN (complex IV inhibitor).

### Lentiviral particles generation and NSCs transduction

To label NSCs, cells were transduced *in vitro* using a third-generation lentiviral carrier (pRRLsinPPT-hCMV) coding for the enhanced farnesylated (f)GFP, which targets the fluorescent protein to the inner plasma membrane of transduced cells [16]. To label mitochondria of NSCs, cells were transduced in vitro using a third generation lentiviral (pLL3.7) carrier coding for the enhanced MitoDsRed expression (i.e. a fusion protein that encodes the leader sequence of cytochrome oxidase IV linked to the florescent protein DsRed), which targets the fluorescent protein to the mitochondrial matrix. The functional stability of these cells (in the absence or in the presence of the lentiviral transcript) has been confirmed with clonal and population studies [54]. Briefly, neurospheres were harvested, dissociated to a single cell suspension and seeded at high density [1.5×10^6^ in a T75 cm^2^ flask (Sigma-Aldrich)] in 5 ml fresh medium. After 12 hrs, 3×10^6^ T.U./ml of lentiviral vectors were added and 6 hrs later additional 5 ml of fresh medium were added. 72 hrs after viral transduction, cells were harvested, re-seeded at normal concentration and transgene expression was measured by FACS analysis.

### L929-Rho^0^ experiments

Mouse Fibroblasts L929-Rho^0^ were a kind gift by Prof Jose’ Antonio Enriquez (CNIC, Madrid Spain). They were grown as adherent cells in fibroblasts medium [DMEM, high glucose, GlutaMAX™ Supplement, pyruvate (Gibco), 10% fetal bovine serum (FBS), 1% pen/strep (Invitrogen) and uridine (Sigma) 50 µg/ml] until they reached confluency (80-90%). The day of passage, cells were washed with PBS (Gibco). Trypsin (0.05% in DMEM) was added at 37°C and inactivated after for 3 min with fibroblast medium (2:1). Cells were collected and spun at 200 x *g* for 5 min, and then re-seeded 1:5 in T175 flasks for normal expansion.

For our experiments, L929-Rho^0^ cells were collected, centrifuged at 200 x *g* for 5 min, then seeded on 13 mm cover slips (5×10^3^ cells/coverslips, BD Bioscience) in 24-wells plates, in 200 μl of uridine supplemented fibroblast medium. The day after, culture media was removed and substituted with selective medium (uridine^-^) or with complete media, according to experimental design. Media change with selective or complete fibroblast medium was carried out every 24h to remove dead cells and/or to supplement with fresh uridine where needed.

After three days upon the switch to selective media, phenotype rescue experiments were carried out by adding EVs or mitochondria obtained from Mito-DsRed^+^ NSCs and resuspended in selective medium (ratios= 1 Rho^0^ L929 fibroblast: EVs/Mito collected from 30 or 90 Mito-DsRed^+^ NSCs). Medium was removed and substituted with selective medium (uridine^-^) or with complete media every 24h until the end of experiment.

We carried out fluorescence stainings and quantifications of uptake of MitoDsRed^+^ particles (3 hrs after treatment) and Rho^0^ L929 survival (5 days after treatment). Rho^0^ L929 fibroblasts were rinsed with PBS and then with PBS with 0.1% Triton X100. To selectively label F-actin, cells were stained with Alexa Fluor 488 phalloidin (1:200, ThermoFisher #A12379), in 0.1% Triton X100 for 20 min at RT and then washed with PBS1X. Nuclei were counterstained with 4’,6-diamidino-2-phenylindole (DAPI) (1:10,000, Invitrogen) for 3 min and eventually coverslips were mounted with Dako mounting kit (Fluka). Uptake of MitoDsRed^+^ particles was quantified with confocal microscopy by counting the number of cells with MitoDsRed^+^ inclusions over the total number of DAPI^+^ cells on 20X objective on 6 regions of interest (ROI) per n≥ 4 replicates per condition.

Rho^0^ L929 survival was quantified on images of the entire coverslip area using an Olympus BX53 microscope with motorized stage and Neurolucida software and a 4X objective. Images were analysed using ImageJ software. Data were represented as number of total DAPI^+^ cells/area.

For Sanger sequencing the DNA from NSCs and Rho^0^ fibroblast L929 was extracted at 16 days after treatment, and the mitochondrial encoded gene *ND3* (*mt-ND3*) was PCR-amplified using KAPA polymerase kit (Biosystems), following manufacturer’s instruction. The primers used were: mt-ND3 forward, 5′-TTCCAATTAGTAGATTCTGAATAAACCCAGAAGAGAGTAAT-3′ and mt-ND3 reverse 5′-CGTCTACCATTCTCAATAAAATTT-3′. PCR amplification protocol was described above. All PCR products were first evaluated on 2% agarose gels (as described above) and then purified using the QIAquick GelExtraction Kit (www.qiagen.com). Sanger sequencing of these products was performed with reverse primers by Source Bioscience (www.sourcebioscience.com). Obtained sequences were visualized and analysed with 4peaks (https://nucleobytes.com/4peaks/index.html).

### Bone marrow derived macrophages (Mφ) culture

Bone marrow derived macrophages (Mφ) were obtained from the bone marrow of C57BL/6 mice, as previously described [49]. Briefly, 9-10 weeks old C57BL/6 female mice were anesthetized with 2% isoflurane and killed by cervical dislocation. Bone marrow was flushed from femurs and tibiae and bone marrow progenitor cells were cultured for 6 days on Petri dishes (Thermo Scientific) in Mφ medium [(DMEM high glucose, Gibco), 10% FBS, 1% pen/strep (Invitrogen) and 10% of conditioned media from L-929 fibroblast cells containing macrophage colony-stimulating factor (mCSF)].

### Analysis of endocytosis/phagocytosis

Mφ were plated in 12 well plates at 150,000 cells per well in 1ml of Mφ medium (DMEM high glucose + 10% dialysed FBS + 1% pen/strep) + 10% mCSF. The cells were let to rest overnight and stimulated with 50 ng/ml lipopolysaccharide (LPS) (Enzo life sciences). After 30 min from the start of the stimulation, cells were treated with either 50 μM dynasore, 20 μM Pitstop 2 (Abcam), or 10 μM cytochalasin D (ThermoFisher). 30 min after drug treatment the cells received either EVs that were extracted from Mito-DsRed^+^ NSCs at a ratio of 1 Mφ : EVs collected from 30 Mito-DsRed^+^ NSCs. Fluorescent labelled 0.04 μm latex beads-Alexa 647 (ThermoFisher) at a concentration of 1:100,000, or fluorescent labelled liposomes-Alexa 647 (Sigma-Aldrich) at a concentration of 0.8 μmoles L-A-phosphatidylcholine, 0.2 μmoles stearylamine and 0.1 μmoles cholesterol per well, were used as controls. After 15 min of treatment, the wells were washed with PBS, fixed with 4% paraformaldehyde (PFA) and stained with F480 and DAPI, before being mounted. Images were taken with a Dragonfly Spinning Disk imaging system (Andor Technologies Ltd.) composed by a Nikon Ti-E microscope, Nikon 60x TIRF ApoPlan and a Zyla sCMOS camera and analysed using the Imaris v.9.1.2 software (Bitplane AG, Zurich, Switzerland). Internalised EVs, latex beads and liposomes were counted from n≥4 ROIs per condition from n= 3 biological replicates. Data were analysed using One-way ANOVA.

### Fluorescent-activated cell sorting (FACS) analysis

Exogenous mitochondria uptake was assessed in LPS-stimulated Mφ at 6hr, following Mito-DsRed^+^ EVs treatment. Mφ were plated in 6 well plates (500,000 cells/well), let rest overnight and stimulated with 50 ng/ml LPS (Enzo life sciences). After 1 hr from LPS stimulation, EVs were extracted from Mito-DsRed^+^ NSCs, resuspended in Mφ medium and added in the same well (ratios: 1 Mφ : EVs collected from 30 Mito-DsRed^+^ NSCs). After 6 hrs, 500 μl of Accumax was added to each well, cells were detached by scraping, washed with 500 μl of PBS 1X and collected. Cells were then centrifuged at 1300 x *g* for 5 min, resuspend in 200 μl of PBS and transferred into FACS tube for the analysis. Subsequently, cells were analysed by flow cytometry using BD LSR Fortessa (BD Biosciences) operated by BD FACS Diva software and at least 100,000 events were collected per sample.

### Quantification of Mito-EVs included in mitochondria, lysosomes and peroxisomes

To analyse the integration of Mito-EVs, Mφ were plated in an 8-well chamber slide (Nunc Lab-Tek II Chamber Slide System, ThermoFisher scientific, Massachusetts, USA) (50,000 cells/well), and left to rest overnight.

For mitochondria staining, Mφ were exposed to 100 nM of MitoTracker Green FM (M-7514, ThermoFisher scientific, Massachusetts, USA) for 10 min and then washed with media. Mφ were then stimulated with 50 ng/ml LPS (Enzo life sciences) and after 1 hr, EVs extracted from Mito-DsRed^+^ NSCs were resuspended in Mφ medium and added in the same well (ratios: 1 Mφ : EVs collected from 30 Mito-DsRed^+^ NSCs). At 6 hrs of incubation, live images and videos (0.15 frames per second, fps) were acquired using a Dragonfly Spinning Disk imaging system (Andor Technologies Ltd.) composed by a Nikon Ti-E microscope, Nikon 60x TIRF ApoPlan and a Ixon EMCCD camera. The videos were processed (8 fps) and the images analysed using the Imaris v.9.1.2 software (Bitplane AG, Zurich, Switzerland). The 3D surface and spots were created for the mitochondrial network and the EVs, respectively. The distance between the surface and spots were calculated using two different thresholds (0 μm to count for including EVs into the network and 0.5 μm for attaching). The analysis of n≥ 10 cells per condition was done blinded (n= 3).

For lysosomes and peroxisomes staining, Mφ were treated as above with LPS and EVs extracted from Mito-DsRed^+^ NSCs. At 6 hrs of incubation cells were washed with PBS three times and fixed with 4% PFA. The anti-LAMP1 (Abcam ab24170,1:500) or PMP70 (Sigma, SAB420018, 1:500) primary antibodies were used to stain for lysosomes or peroxisomes. Images were acquired using a Dragonfly Spinning Disk system as described above but using a Zyla sCMOS camera instead. MitoDsRed^+^ EVs co-localisation analysis was performed in n≥ 10 cells per condition using Imaris software and X coefficient was calculated (n= 3).

### Quantification of mitochondrial network morphology

Mφ were plated on coverslips (Microscope Cover Glasses, Glaswarenfabrik Karl Hecht GmbH, Sondheim, Germany) at 50,000 cells/well and the next day stimulated with 50 ng/ml LPS (Enzo life sciences). After 1 hr of LPS stimulation, Mφ were treated with or without EVs extracted from Mito-DsRed^+^ NSCs for another 6 hrs, washed with PBS three times and fixed with 4% PFA. Cells were then permeabilised with 0.1% Triton X-100, blocked in 5% goat serum and stained with an anti-TOMM20 antibody (rabbit, Santa Cruz, Dallas, USA, 1:300). Representative images were acquired as described above for the integration analysis. The mitochondrial network analysis was performed in a blind way scoring each cell (n = 3) as fused, intermediated or fragmented in respect of their mitochondrial network (average of n = 140 cells/experiment).

### Extracellular flux (XF) assays

A XF24e Extracellular Flux Analyser (Seahorse Bioscience) was used for all XF assays.

Mφ were seeded 6 days after bone marrow isolation with fresh Mφ medium on a 24 well XF24 cell culture microplate (1×10^5^ cells/well). After approximately 18 hrs from seeding, Mφ were stimulated by adding 50 ng/ml LPS (Enzo life sciences). After 1h from LPS stimulation, EVs were extracted from NSCs, resuspended in Mφ medium and added in the same well (ratios: 1 Mφ : EVs collected from 30 NSCs). At 6 hrs from treatment, Mφ medium was replaced with XF medium [Seahorse salt solution (Seahorse Bioscience), 1% glutamine 200 mM, 1% pyruvate 100 mM, 1% FBS, D-glucose (225 mg/50ml final volume)] pH 7.35-7.45, and baseline oxygen consumption rate (OCR) and extracellular acidification rate (ECAR) were measured for 3 reads. Mitochondrial stress protocol was performed using oligomycin, carbonyl cyanide-4-(trifluoromethoxy)phenylhydrazone (FCCP), rotenone and antimycin following manufacturer’s instructions (XF cell mito stress test kit, Seahorse Bioscience).

For experiments in which EVs were pre-exposed to the uncoupling agent FCCP, EVs were isolated as previously described. After the ultracentrifugation step at 100,000 x *g* for 70 min at 4°C, EVs were resuspended in 1 ml of Mφ medium with 1μM FCCP for 1 hr at 37 °C (EVs^FCCP^), or Mφ medium alone (EVs). Particles were then washed with a centrifugation step at 100,000 x g for 30 min at 4°C, followed by resuspension in 200 μl of PBS and an additional wash with 3.2 ml of PBS. Tubes were then subjected to an additional centrifugation at 100,000 x *g* for 30 min at 4°C using an optima MAX ultra with TLA110 fixed angle rotor, the supernatant was completely discarded and EVs^FCCP^ (or EVs) were finally resuspended in Mφ medium at the desired concentration.

After the completion of each XF assay, cells were washed with PBS and 25 µl of 1X RIPA buffer (with protease/phosphatase inhibitors) were added to each well. The total protein amount/well was estimated with a BCA Protein Assay Kit (Thermo Scientific) and used to normalize the OCR and ECAR values of the single well.

### Quantitative gene expression analysis (qPCR) and microarrays

Experiments in which EVs were pre-exposed to the uncoupling agent FCCP were performed as reported above.

For quantitative gene expression analysis (qPCR) total RNA was extracted using RNeasy kit (Qiagen) according to manufacturer’s instructions. Total RNA (400 ng) was retro-transcribed into complementary DNA (cDNA) using cDNA Reverse Transcription kit (Life Technologies) according to the manufacturer’s instructions. qPCR was performed using TaqMan Universal PCR Master Mix (Applied Biosystems) and TaqMan Gene Expression Assays for: *Il1* (Mm00434228_m1, Life Technologies), *Nos2* (Mm00440502_m1, Life Technologies) and *Il6* (Mm00446190_m1, Life Technologies). *Actb* (ACTB, REF4351315 Life Technologies) was used as internal calibrator. Samples were tested in triplicate on a Quant StudioTM 7 Flex Real-Time PCR System (Applied Biosystems) and analysed using the 2^-ΔΔCT^ method. Briefly, the threshold cycle (CT) method uses the formula 2^-ΔΔCT^ to calculate the expression of target genes normalized to a calibrator. The CT indicates the cycle number at which the amount of amplified target reaches a fixed threshold. The CT values were normalized for endogenous reference [ΔCT = CT (target gene) – CT (*Actb*, endogenous reference)] and compared with a calibrator using the ΔΔCT formula [ΔΔCT = ΔCT (sample) – ΔCT (calibrator)].

For microarrays, samples were prepared according to Affymetrix protocols (Affymetrix, Santa Clara, CA). RNA quality and quantity were ensured using the Bioanalyzer (Agilent, Santa Clara, CA) and NanoDrop (Thermo Scientific, Waltham, MA) respectively. For RNA labelling, 200 ng of total RNA was used in conjunction with the Affymetrix recommended protocol for the Clariom_S chips. The hybridization cocktail containing the fragmented and labelled cDNAs was hybridized to the Affymetrix Mouse Clariom_S GeneChip. The chips were washed and stained by the Affymetrix Fluidics Station using the standard format and protocols as described by Affymetrix. The probe arrays were stained with streptavidin phycoerythrin solution (Molecular Probes, Carlsbad, CA) and enhanced by using an antibody solution containing 0.5 mg/mL of biotinylated anti-streptavidin (Vector Laboratories, Burlingame, CA). An Affymetrix Gene Chip Scanner 3000 was used to scan the probe arrays. Gene expression intensities were calculated using Affymetrix AGCC software. Downstream analysis was conducted in R/Bioconductor [25].

The annotation package for the Clariom_S chips was retrieved from Bioconductor. The CEL files were then loaded into R, RMA normalized with the oligo package, and filtered to only retain probes annotated as “main” [8]. Differential expression testing was performed using limma and the resulting p.values were corrected with the Benjamini-Hochberg method. Kyoto Encyclopedia of Genes and Genomes (KEGG) pathway enrichment analyses were performed using the Generally Applicable Gene-set Enrichment (GAGE) package with default parameters [39]. Pathways of interest were visualised with Pathview [38]. The microarray raw data were deposited in ArrayExpress with the accession number E-MTAB-8250.

### EAE induction, treatment and behavioural studies

Animal research has been regulated under the Animals (Scientific Procedures) Act 1986 Amendment Regulations 2012 following ethical review by the University of Cambridge Animal Welfare and Ethical Review Body (AWERB). Animal work was covered by the PPL 80/2457 (to SP).

C57BL/6 female mice were immunised with myelin oligodendrocyte glycoprotein (MOG)-induced experimental autoimmune encephalomyelitis (EAE), as previously described [49]. Briefly, mice were anaesthetized with isoflurane (4% induction, 1.5% maintenance), and received n= 3 subcutaneous (s.c.) injections (2 flanks and 1 at the base of the tail) of 50 μl containing 200 μg/mouse MOG35-55 (Multiple Peptide System) (Espikem), incomplete Freund’s Adjuvant (IFA) and 8 mg/ml *Mycobacterium tuberculosis* (Scientific Laboratories Supply). 100 μl of Pertussis Toxin (5 ng/μl) (List Biological Laboratories) was injected intravenously (i.v.) on the day of the immunization and again after 48 hrs.

Body weight and EAE clinical score (0 = healthy; 1 = limp tail; 2 = ataxia and/or paresis of hindlimbs; 3 = paralysis of hindlimbs and/or paresis of forelimbs; 4 = tetraparalysis; 5 = moribund or death) were recorded daily.

After 11-19 days post immunisation (dpi), mice developed the first clinical signs of diseases (disease onset). At 3 days after disease onset (= peak of disease, PD), mice with similar scores were randomly assigned to the different treatment groups. After randomization, mice received a single intracerebroventricular (ICV) injection (AP -0.15, ML +1.0 left, DV -2.4) of fGFP^+^/MitoDsRed^+^ NSCs (1×10^6^ in 5 μl PBS) or EVs derived *in vitro* from fGFP^+^/MitoDsRed^+^ NSCs (64μg in 5 μl PBS). EAE mice injected ICV with 5 μl PBS were used as controls.

### *Ex vivo* tissue pathology

At 55 dpi, mice were deeply anesthetized with an intraperitoneal (i.p.) injection of ketamine 10 mg/ml (Boehringer Ingelheim) and xylazine 1.17 mg/ml (Bayer) in sterile water and transcardially perfused with 1ml EDTA 5M in 500ml saline 0.9% NaCl for 5 min, followed by a solution of 4% PFA in PBS for another 5 min.

Brains and spinal cords were isolated and post-fixed in 4% PFA in PBS at 4°C overnight. Tissues were then washed in PBS and left for at least 48-72 hrs in 30% sucrose in PBS at 4°C for cryo-protection. Brains and spinal cords were then embedded in optimum cutting temperature (OCT) medium, frozen with liquid nitrogen and cryo-sectioned (25 μm coronal section thickness for brains and 10 μm axial section thickness for spinal cords) using a cryostat (CM1850, Leica, Wetzlar, Germany) with a microtome blade (A35, Feather, Osaka, Japan). Sections were then stored at - 80°C until use.

For quantification of mitochondrial transfer events, sections were rinsed with PBS, and then blocked for 1 hr at RT in blocking buffer (0.1% Triton X100 and 10% secondary antibody species serum in PBS). A Fab fragment affinity purified IgG anti-mouse was applied if anti-mouse antibodies were used (1:10, Jackson ImmunoResearch). The following primary antibodies, diluted in blocking buffer, were used at 4°C overnight: anti-GFP (1:250, Invitrogen), anti-DsRed(RFP) (1:400, Abcam), anti-GFAP (1:500, Abcam), anti-NeuN (1:250, Chemicon), anti-Olig2 (1:500, Millipore), anti-CD3 (1:250, Abcam) or anti-F4/80 (1:100 Serotec). Sections were then washed in PBS with 0.1% Triton X100 and incubated with the appropriate fluorescent secondary antibodies (1:1,000 Alexa-fluor 405, 488, 555, 647, Invitrogen) for 1 hr at RT. After washing in PBS, nuclei were counterstained with DAPI (1:10,000, Invitrogen) for 3 min and then mounted with Dako mounting kit (Fluka). Nonspecific staining was observed in control incubations in which the primary antibodies were omitted.

Quantification of mitochondria transfer events was obtained from randomized n≥ 12 brain ROIs acquired using a confocal microscope (Leica TCS SP5 Microscope) and data are expressed as % ± SEM of fGFP^-^/MitoDsRed^+^ particles over total fGFP^-^/MitoDsRed^+^ particles found in GFAP^+^ astrocytes, NeuN^+^ neurons, Olig2^+^ oligodendrocytes, CD3^+^ T cells or F4/80^+^ mononuclear phagocytes (an average of 292.5 total fGFP^-^/MitoDsRed^+^ particles per cell type were quantified).

### Statistical analysis

Statistical analyses of all data were performed with Graph Pad Prism (version 8.00 for Mac, GraphPad Software). Differences among groups were analysed using One-Way-ANOVA followed by a Tukey’s multiple comparison test (unless otherwise stated). Values are given in the text and figures as mean values ± SEM and a p value < 0.05 was accepted as significant in all analyses (unless otherwise stated).

## Results

### Proteomic analysis of NSC-derived EVs and exosomes identifies mitochondrial proteins

We first performed an untargeted multiplex Tandem Mass Tag (TMT)-based proteomic analysis of the whole EV fraction and sucrose-gradient purified exosomes spontaneously released by NSCs *in vitro* and compared them with parental NSC whole-cell lysates (Fig. 1a and Table S1).

**Fig. 1.**
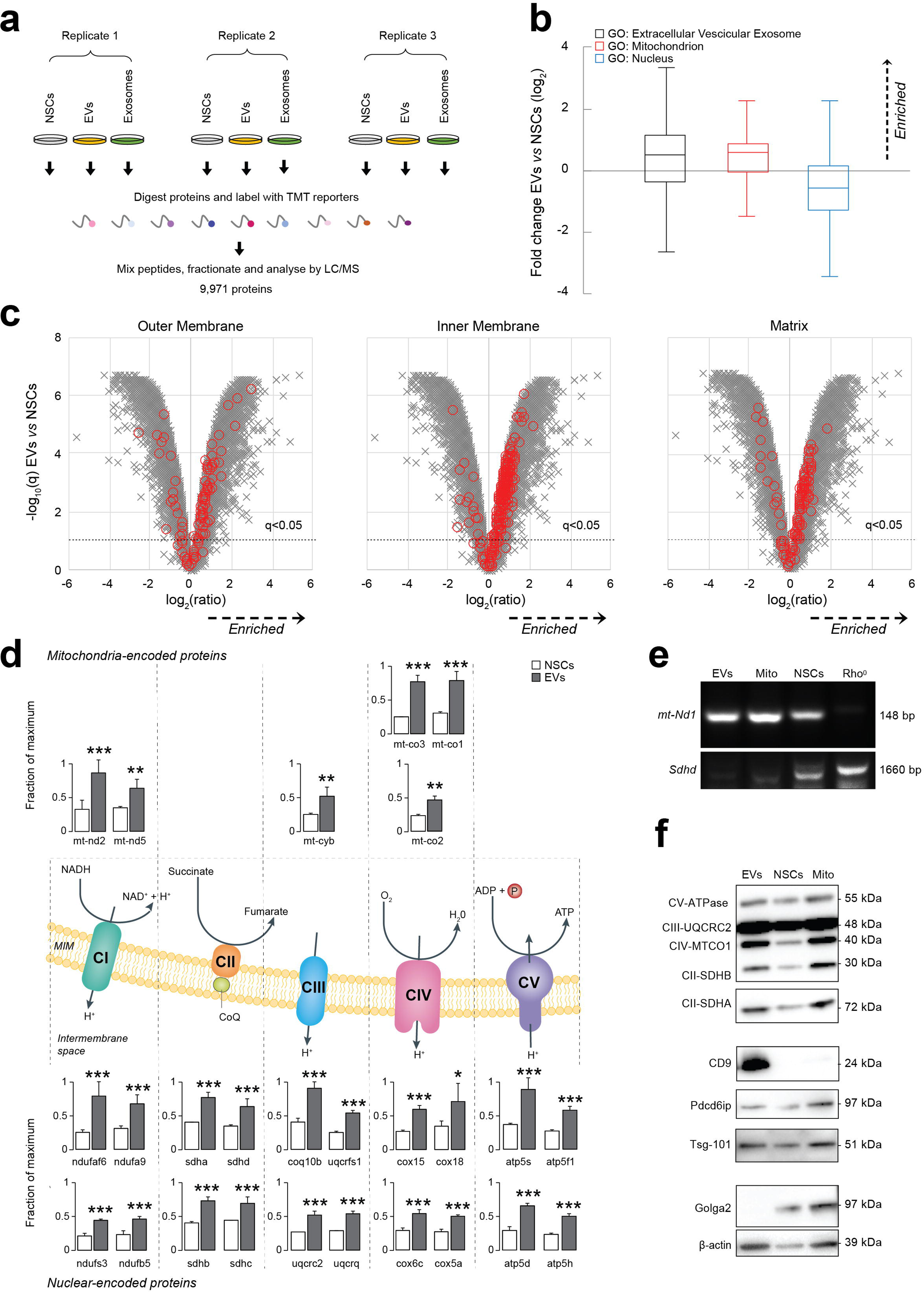
NSCs spontaneously release mitochondrial proteins and mtDNA. **a**, Overview of multiplex TMT-based proteomic experiment. TMT-based proteomics identified a total of 9,971 proteins, of which 9,951 were quantitated across all conditions. **b**, Relative abundance of proteins annotated with indicated Gene Ontology Cellular Component (GOCC) subcellular localisations in extracellular vesicles (EVs) compared with NSC whole cell lysates (NSCs). Annotations were available for 9,049/9,971 cellular proteins identified in the multiplex TMT-based functional proteomic experiment illustrated in a. Boxplots show median, interquartile range and Tukey whiskers for proteins with the following annotations: extracellular vesicular exosome (GO:0070062, black outline, enriched in EVs); mitochondrion (GO:0005739, red outline, enriched in EVs); nucleus (GO:0005634, blue outline, depleted from EVs). Data are from 3 independent biological replicates. **c**, Relative abundance of proteins from different mitochondrial compartments (outer membrane, matrix and inner membrane) in EVs compared with NSCs. Volcano plots show statistical significance (y axis) *vs* fold change (x axis) for 9,951/9,971 cellular proteins quantitated across all 3 biological replicates (no missing values) in the multiplex TMT-based functional proteomic experiment illustrated in a. Proteins annotated with the following GOCC subcellular localisations are highlighted in red: mitochondrial outer membrane (GO:0005741, enriched in EVs); mitochondrial inner membrane (GO: 0005743, enriched in EVs); mitochondrial matrix (GO:0005759, enriched in EVs). A false discovery rate (FDR) threshold of 5% is indicated (proteins with Benjamini-Hochberg FDR-adjusted p values (q values) < 0.05). **d,** Relative abundance of selected mitochondrial proteins in EVs compared with NSCs. Mitochondrial complex (C) proteins enriched in EVs and encoded in the mitochondrial (upper panel) or nuclear (lower panel) genomes in the multiplex TMT-based functional proteomic experiment include: NADH:ubiquinone oxidoreductase or CI [mtnd2 (ND2 subunit), mtnd5 (ND5 subunit), ndufaf6 (assembly factor 6), ndufa9 (subunit A9), ndufs3 (core subunit S3), ndufb5 (1 beta subcomplex subunit 5)], succinate dehydrogenase or CII [Sdha (Subunit A), sdhd (cytochrome b small subunit), sdhb (iron-sulfur subunit), sdhc (cytochrome b560 subunit)], cytochrome b-c1 or CIII [mt-cyb (cytochrome B), Uqcrfs1 (subunit 5), Uqcrc2 (subunit 2), coq10b (coenzyme Q10B), uqcrq (subunit 8)], cytochrome C oxidase or CIV [mt-co3 (oxidase III), mtco1 (oxidase I), mtco2 (oxidase II), cox15 (subunit 15), cox18 (assembly protein 18), cox6c (subunit 6C), cox5a (subunit 5a)] and ATP synthase or CV [atp5f1 (subunit gamma), atp5s (subunit S), atp5d (subunit delta), atp5h (subunit D)]. Mean abundances (fraction of maximum) and 95% confidence intervals (CIs) from 3 biological replicates are shown. *q < 0.05, **q < 0.01, ***q < 0.001. **e,** Representative PCR amplification of DNA extracted from NSCs, EVs and isolated mitochondria (Mito). The mitochondrial encoded gene *mt-ND1* (NADH-ubiquinone oxidoreductase chain 1) was found to be present in EVs, Mito and NSCs (L929 Rho^0^ were used as negative controls). **f**, Representative protein expression analysis by western blot of NSCs, EVs and isolated mitochondria (Mito). Mitochondrial complex proteins (CV-ATPase, CII-SDHA, CII-SDHB, CIV-MTCO1, CIII-UQCRC2), EV positive markers (Tsg-101, Pdcd6ip, CD9) and EV negative markers (Golga2) are shown, as well as β-actin.

Using Gene Ontology Cellular Component (GOCC) annotations, we investigated the subcellular origin of proteins enriched in EVs compared with NSCs (Fig. 1b). We found that proteins with annotations indicating exosomal localisation were markedly enriched in EVs, whereas proteins with annotations indicating nuclear localisation were depleted (Fig. 1c). Interestingly, proteins with annotations indicating mitochondrial localisation were also significantly enriched in EVs *vs* NSCs (Fig. 1b).

To further investigate this finding, we next examined our data with more specific GOCC daughter annotations indicating localisation to the three major mitochondrial structural components: outer membrane, inner membrane and matrix. We found that proteins with these annotations were relatively enriched in EVs compared to parental NSCs (Fig. 1c). We also specifically scrutinised the relative abundances of subunits of the five mitochondrial complexes and discovered that proteins coded in both the mitochondrial and nuclear genomes were all significantly enriched in EVs (Fig. 1d). When we investigated specifically the DNA content of the EVs, we found that the mitochondrial gene NADH dehydrogenase subunit 1 (*mt-ND1*), which is encoded in the mtDNA, was present in EVs. However, this was not the case for the mitochondrial gene succinate dehydrogenase complex subunit D (*Sdhd*), which is encoded in the nuclear DNA (Fig. 1e), thus showing that nuclear DNA was not enriched but instead suggesting the likely presence of mitochondria with intact mitochondrial matrix in EVs.

To further validate our TMT-based proteomic data using an orthogonal technique, we next subjected EVs and NSCs to immunoblot analysis. EVs were enriched in exosomal markers (CD9, Pdcd6ip, Tsg101) and mitochondrial complexes, but depleted of Golgi markers (Golga2) (Fig. 1f) compared to NSCs. Conversely, a control preparation enriched of mitochondria (Mito) isolated from NSCs [15] was found to be depleted of CD9 and enriched in Golga2.

To exclude any potential bias related to our own purification methods, we further employed two additional high-quality and scalable exosome/EV isolation protocols that avoid ultracentrifugation [21, 29] (Fig. 2a). We found that all protocols yielded EVs depleted of Golga2 but enriched in CD9 and mitochondrial proteins. In addition, we also tested another ultracentrifugation protocol that adds an additional 0.22 μm ultrafiltration step [68] and were able to readily detect mitochondrial complexes (Fig. 2b). Mitochondrial proteins are therefore found in NSC EVs, irrespective of the protocols used to isolate EVs from tissue culture media.

**Fig. 2.**
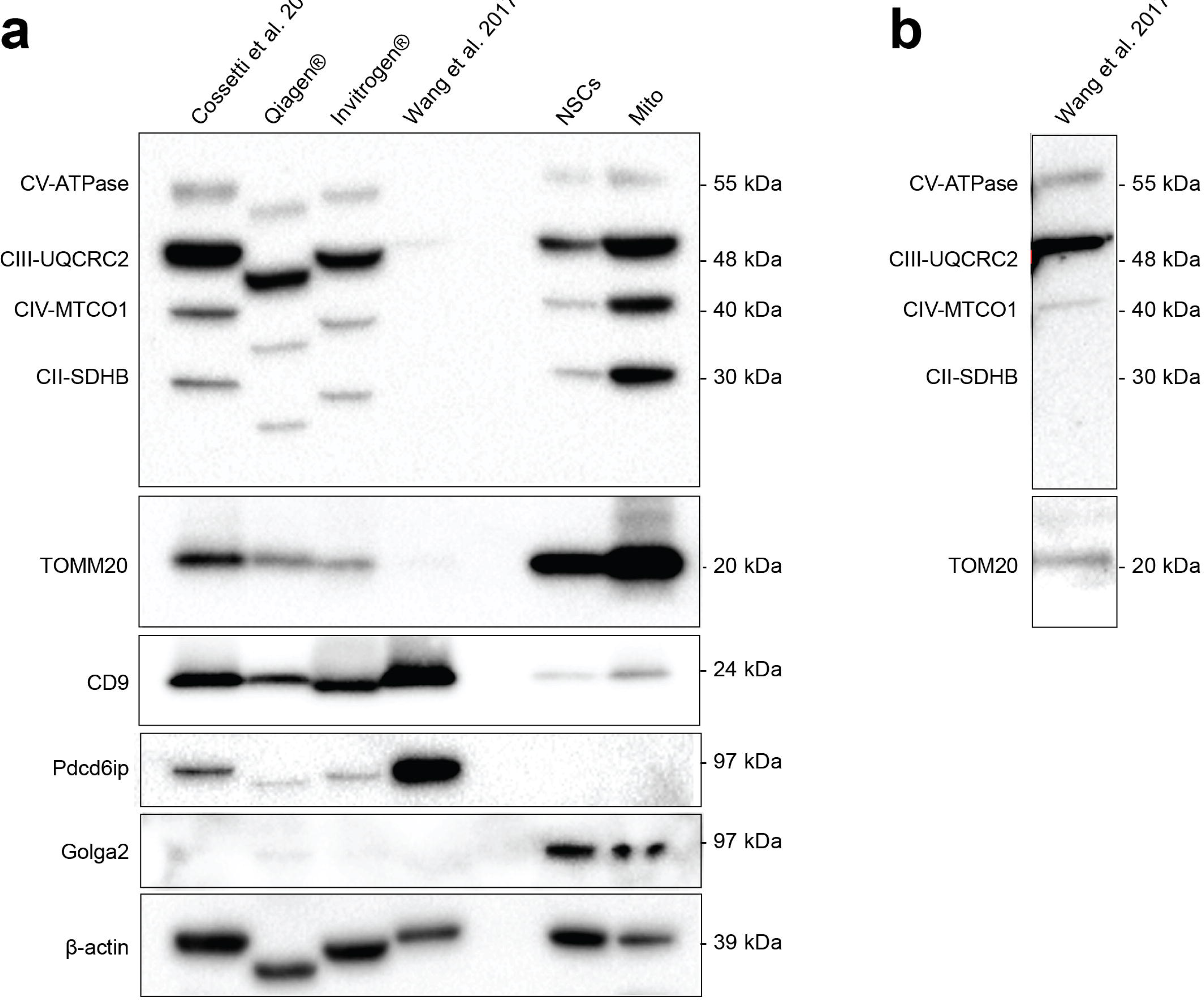
NSC-derived EVs isolated with alternative protocols contain mitochondrial proteins. **a,** Protein expression by western blot analysis of EVs isolated using in house protocol, with commercial kits (Qiagen cat. No 76743 and Invitrogen cat. No 4478359), and an alternative protocol with an additional 0.22 μm ultrafiltration step. NSCs and isolated mitochondria (Mito) are used as comparative controls. Mitochondrial complexes proteins (CV-ATPase, CII-SDHA, CII-SDHB, CIV-MTCO1, CIII-UQCRC2) mitochondrial outer membrane translocase (TOMM20), exosomal positive (Pdcd6ip, CD9) and negative (Golga2) markers are shown, as well as β-actin. **b**, Longer exposure of the lane containing the EVs isolated with the alternative protocol with an additional ultrafiltration step showing mitochondrial proteins.

We next focused on the exosomal fraction isolated via sucrose gradient fractionation from the EV preparation, as described [9]. Compared with parental NSCs, the overall protein composition of exosomes by TMT-based proteomic analysis was similar to EVs (Table S1). However, we found that numerous proteins were selectively depleted in exosomes by the additional purification step compared to EVs (Fig. 3a). Exosomes were also significantly enriched in proteins with GOCC annotations indicating localisation to the mitochondrial outer membrane, inner membrane, and matrix compared to parental NSCs (Fig. 3b). Immunoblot analysis confirmed that fractions 6-9 corresponding to the expected exosomal density were all enriched in mitochondrial complex proteins (Fig. 3c). When looking at the DNA content of the single fractions, we identified the mitochondrial gene *mt-ND1* but not the *Sdhd* gene (Fig. 3d), which unambiguously confirms that NSC exosomes, as well as EVs, harbour mitochondrial proteins and mtDNA.

**Fig. 3.**
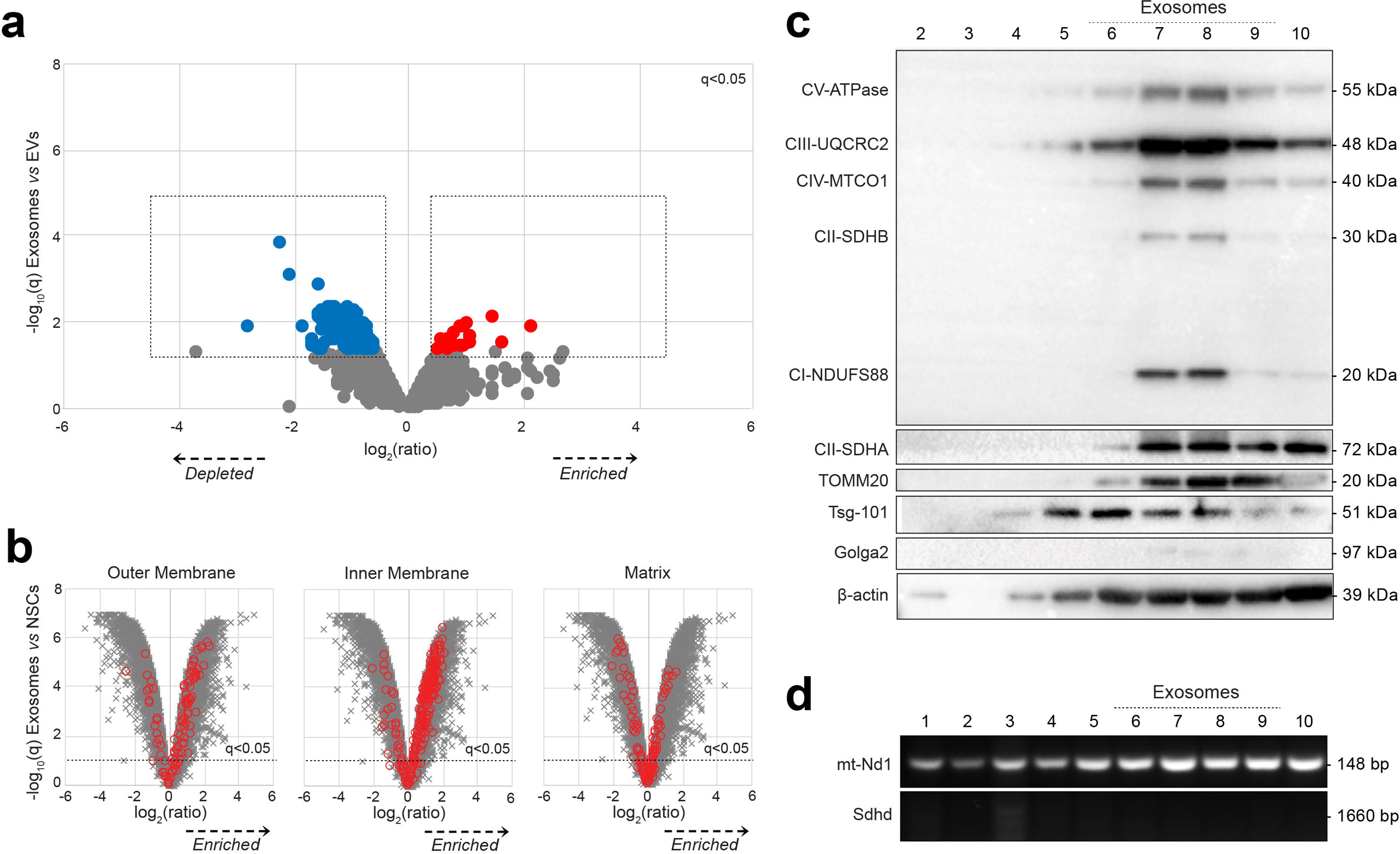
Proteomic analysis of EVs fractions obtained via sucrose gradient. **a**, Relative abundance of proteins in exosomes (fractions 6-9) compared with EVs. Volcano plot shows statistical significance (y axis) *vs* fold change (x axis) for 9,951/9,971 cellular proteins quantitated across all 3 biological replicates (no missing values) in the multiplex TMT-based functional proteomic experiment illustrated in Fig. 1a. A total of 187 proteins were found to be significantly depleted in exosomes, *vs* EVs (blue), while 25 proteins were significantly enriched (red); q < 0.05 (Table S1). **b**, Relative abundance of proteins from different mitochondrial compartments (outer membrane, matrix and inner membrane) in exosomes (fractions 6-9) compared with NSCs. Volcano plots show statistical significance (y axis) *vs* fold change (x axis) for 9,951/9,971 cellular proteins quantitated across all 3 biological replicates (no missing values) in the multiplex TMT-based functional proteomic experiment illustrated in Fig. 1a. Proteins annotated with the following GOCC subcellular localisations are highlighted in red: mitochondrial outer membrane (GO:0005741, enriched in EVs); mitochondrial matrix (GO:0005759, enriched in EVs); mitochondrial inner membrane (GO: 0005743, enriched in EVs). A false discovery rate (FDR) threshold of 5% is indicated (proteins with Benjamini-Hochberg FDR-adjusted p values (q values) <0.05). **c**, Representative protein expression analysis by western blot of EVs fractions (2-10) obtained via continuous sucrose gradient. Fractions 6-9 (corresponding to the expected exosomal density between 1.13 and 1.21 g/ml) were specifically enriched for mitochondrial complex proteins (CV-ATPase, CII-SDHA, CI-NDUFS88, CII-SDHB, CIV-MTCO1, CIII-UQCRC2), for the mitochondrial outer membrane translocase TOMM20 and the exosomal marker Tsg-101 (while they were negative for the Golgi marker Golga2). β-actin is also shown. **d**, Representative PCR amplification of DNA extracted from EVs fractions (2-10) obtained via continuous sucrose gradient. The mitochondrial encoded gene *mt-ND1* (NADH-ubiquinone oxidoreductase chain 1) was found to be present in most of the EVs fractions, while the nuclear encoded mitochondrial gene *Sdhd* (succinate dehydrogenase complex subunit D) was were used as negative control.

### Structurally and functionally intact mitochondria are found in NSCs derived EVs preparations

We next characterised the whole EV fraction released *in vitro* by NSCs using tunable resistive pulse sensing (TRPS) analysis and nanoparticle tracking analysis (NTA).

We found that EVs had a mode diameter ranging between 80 nm and 150.8 nm respectively, as described [9] (Fig. S1). This size distribution was then further investigated with a morphological analysis based on transmission electron microscopy (TEM). Amongst a heterogeneous population of EVs with a mode diameter of 286.7 nm, we identified several mitochondria-like structures, either free or encapsulated in other membranes, with a mean diameter of 695.4 nm (± 68.47 nm) (Fig. 4a). We next used cryo-transmission electron microscopy (Cryo-TEM) [2] combined with immuno-gold labelling using an anti-TOMM20 antibody conjugated to gold nanoparticles to further confirm this finding. TOMM20 positive mitochondria were found in all EV preparations (Fig. 4b). Altogether these complementary approaches suggest that *in vitro* NSCs spontaneously release EVs in the sub-micron range (<1000 nm) that include structurally intact mitochondria (Mito-EVs).

**Fig. 4.**
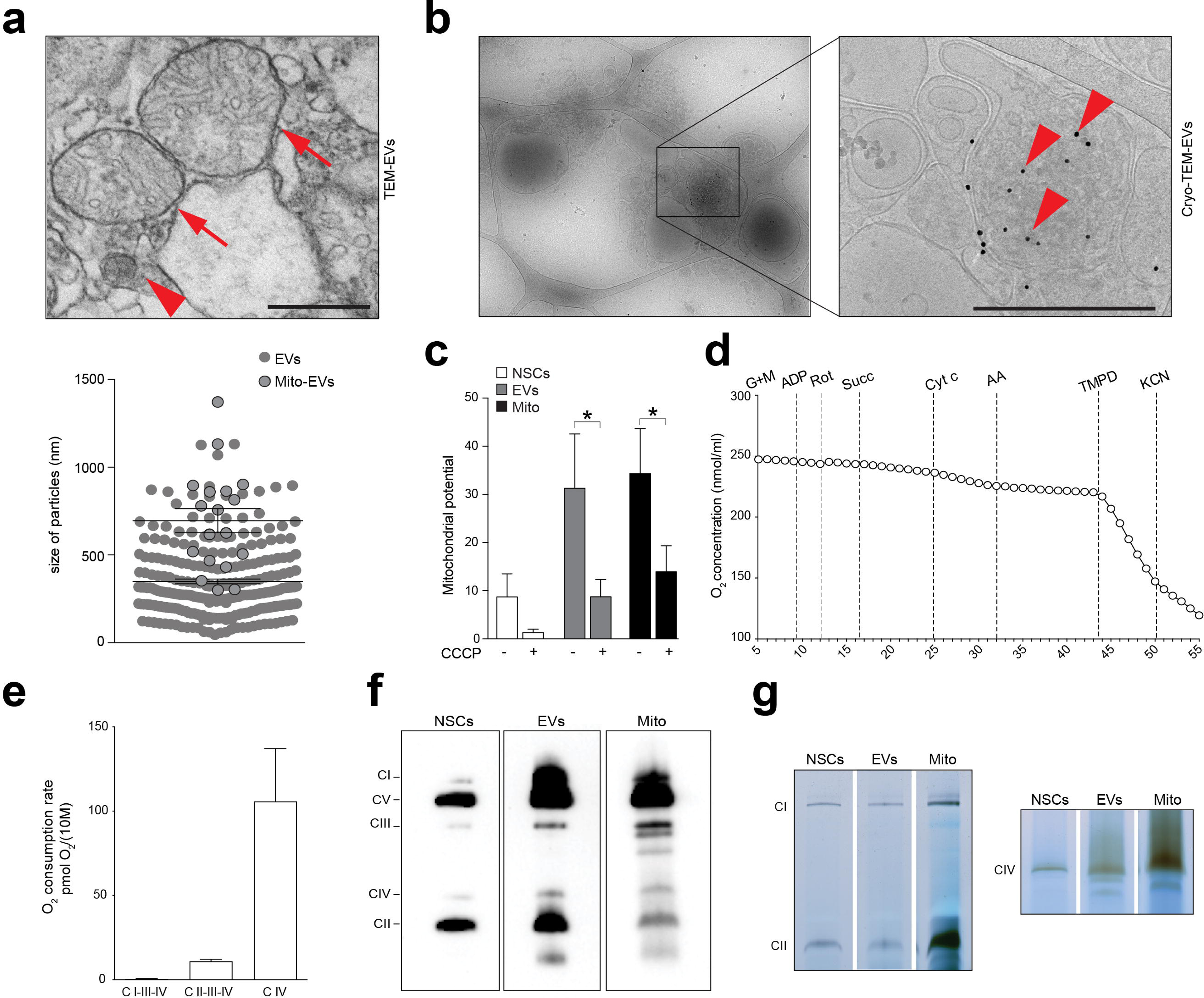
NSCs-derived EV preparations contain structurally and functionally intact mitochondria. **a**, Representative Transmission Electron Microscopy (TEM) image of NSCs-derived EVs showing free (arrows) and encapsulated (arrowhead) mitochondria. Scale bar: 500 nm. The graph shows size particle analysis and quantification of mitochondria (black circled grey dots) found in the EV preparations (grey dots). Data are expressed as mean values (± SEM) from 3 replicates. **b**, Representative cryo-transmission electron microscopy (Cryo-TEM) image of NSC-derived EVs labelled with anti-TOMM20-gold nanoparticles (NP) antibodies conjugated to gold NP, demonstrating the presence of intact mitochondria. Scale bar: 500 nm. **c**, Mitochondrial membrane potential of NSCs, EVs and Mito preparations treated (or not) with the mitochondrial uncoupler carbonyl cyanide m-chlorophenyl hydrazone (CCCP). *p≤ 0.05. Data are expressed as mean JC1 Ratio (red/green fluorescence signal) (± SEM) per unit (unit defined as 10^6^ NSCs and the EVs/Mito produced by 10^6^ NSCs) from n ≥ 4 experiments. **d**, Mitochondrial respiration of permealized EVs detected by high resolution respirometry (HRR). Representative plot showing O_2_ concentration changes over time upon serial additions of selected mitochondrial complexes substrates, inhibitors and uncouplers, including CI substrates (glutamate and malate: G+M, adenosine diphosphate: ADP), CI inhibitor (rotenone: Rot), CII substrate (succinate: Succ), cytochrome *c* (Cyt c) to compensate for a possible loss due to outer membrane disruption, CIII inhibitor (antimycin A: AA), CIV electron donor (*N, N, N′, N′*-tetramethyl-p-phenylenediamine: TMPD) and CIV inhibitor (potassium cyanide: KCN). **e**, Complex respiratory rate in EVs. Data are mean values (± SEM) from n = 3 independent experiments and shown as oxygen consumption rate pmolO_2_/ normalized by the EVs produced by 10^6^ NSCs. **f**, Representative image of mitochondria respiratory chain native complexes separated by Blue Native polyacrylamide gel electrophoresis (BN-PAGE) showing the presence of structurally intact respiratory complexes (CI-V) in NSCs, released EVs, and isolated Mito preparations. **g**, Representative image of in situ gel activity of CI-II-IV in NSCs, EVs and Mito obtained from BN-PAGE gel incubation for 24h.

We then tested the functionality of these Mito-EVs by analysing the activity of the electron transport chain (ETC) in maintaining a mitochondrial transmembrane potential and respiration, using a JC1 assay and high resolution respirometry (HRR) [42], respectively. EVs indeed showed a conserved mitochondrial membrane potential, which was responsive to the mitochondrial uncoupler carbonyl cyanide *m*-chlorophenyl hydrazone (CCCP) (Fig. 4c). In addition, EVs exhibited oxygen consumption when the substrates for the mitochondrial complexes were added to the EV preparation (Fig. 4d,e). When we tested the activity of the mitochondrial respiratory chain complexes, using blue native polyacrylamide gel electrophoresis (BN-PAGE) [47], intact protein complexes were isolated from EVs in native conditions (Fig. 4f). This finding was coupled with a conserved catalytic activity of CI-CII-CIV, which suggests the presence of functionally intact mitochondrial complexes (Fig. 4g) [70].

Altogether these findings demonstrate that Mito-EVs released by NSCs harbour a functional mitochondrial ETC and the potential for oxidative phosphorylation (OXPHOS).

### Mito-EVs revert the auxotrophy of mtDNA deficient cells

We next investigated whether NSC Mito-EVs have any effect on target cells. To this end, we generated NSCs that constitutively express the mitochondrial Mito-DsRed fluorescent reporter (MitoDsRed^+^ NSCs) to stably label Mito-EVs. We then used Mito-DsRed^+^ NSCs to derive EVs (MitoDsRed^+^-EVs) and treat cells that had been depleted of mtDNA using extended low-dose ethidium-bromide treatment [62] (Fig. 5a). These L929 Rho^0^ have an auxotrophic growth and dependence on extracellular uridine, which satisfies their energy demand despite the inhibition of dihydroorotate dehydrogenase (DHODH) allowing for cell survival [14].

**Fig. 5.**
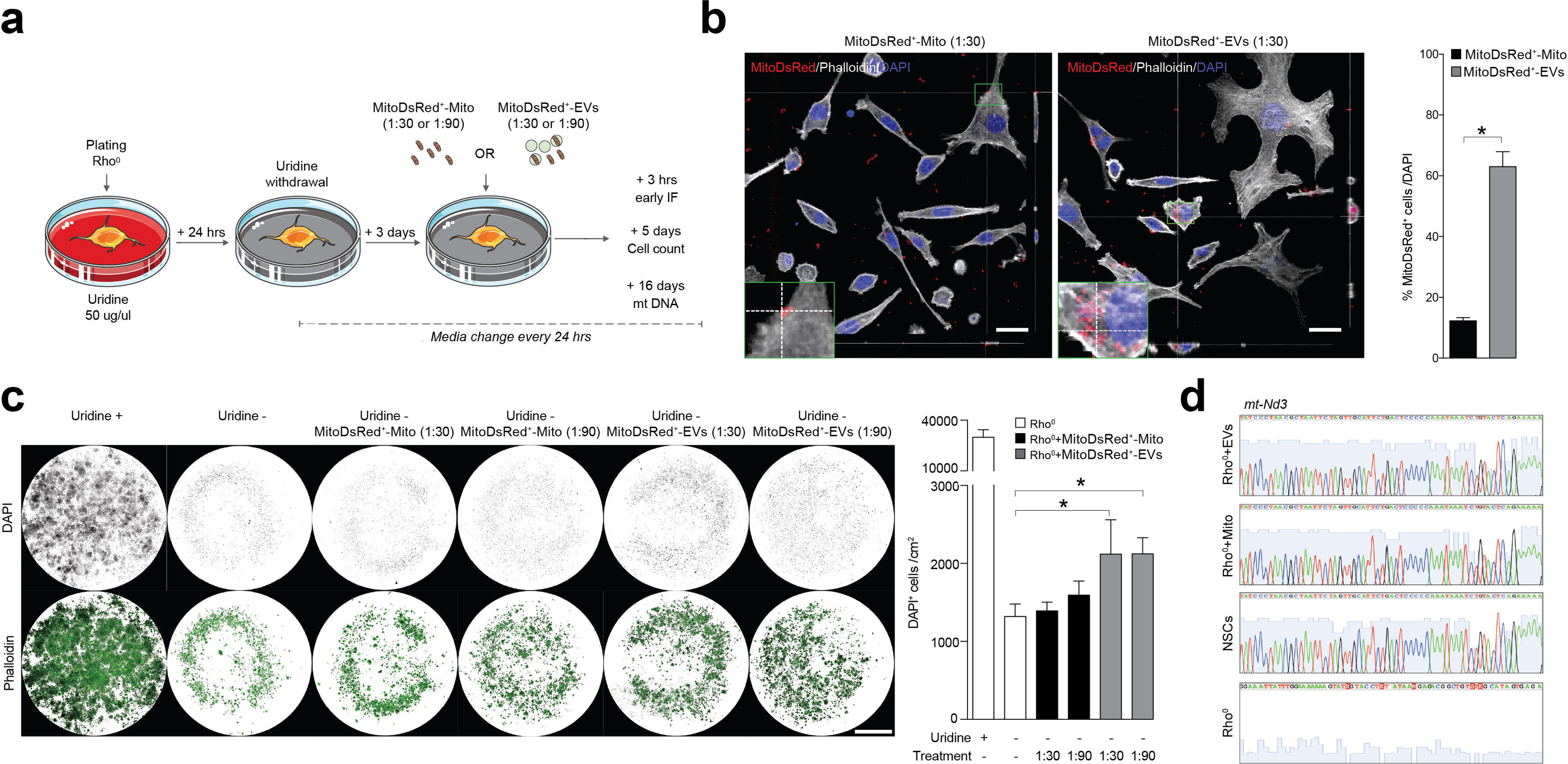
NSC-derived EVs can correct mitochondrial DNA deficiency of L929 Rho^0^ cells. **a**, Experimental setup for *in vitro* studies with L929 Rho^0^ cells. L929 Rho^0^ cells were deprived of uridine (Uridine^-^) and then treated with either MitoDsRed^+^ EVs (ratio 1:30 or 1:90) or MitoDsRed^+^ Mito (ratio 1:30 or 1:90) after 3 days. **b**, Representative confocal images and quantification of Uridine^-^ L929 Rho^0^ cells showing incorporation of MitoDsRed^+^ mitochondria at 24h from treatment with either EVs or Mito (ratio 1:30). Orthogonal section (XY) of Z-stacks is shown. Data are mean percentage over total DAPI^+^ cells (± SEM) from n = 9 cells per replicate, n = 2 replicates per condition). Scale bars: 25 μm. **c**, Representative images and quantification of L929 Rho^0^ cells surviving 5 days after treatment with either EVs or Mito. Data are number of DAPI^+^ cells/cm^2^ (± SEM) from 4 replicates per condition. *p ≤ 0.05. Scale bars: 3.25 mm. **d**, Sanger sequencing chromatograms showing the mitochondrial encoded gene *mt-ND3* in L929 Rho^0^ cells at 16 days after treatment with either EVs or Mito. NSCs and L929 Rho^0^ cells are used as positive and negative controls, respectively.

In conditions of uridine deprivation, we found that L929 Rho^0^ cells efficiently incorporated MitoDsRed^+^-EVs within 24 hrs from treatment compared to L929 Rho^0^ cells treated with a preparation enriched of isolated mitochondria (MitoDsRed^+^-Mito) (Fig. 5b). At 5 days post treatment, while only a minority of untreated L929 Rho^0^ cells survived uridine depletion, L929 Rho^0^ cells treated with MitoDsRed^+^-EVs displayed a significantly higher survival (Fig. 5c). Finally, at 16 days post treatment, we were able to sequence the *mt-ND3* mitochondrial gene from L929 Rho^0^ cells treated with MitoDsRed^+^-EVs (Fig. 5d), which indicates the efficient integration of (exogenous) Mito-EV-derived mtDNA in target cells and a correction of their intrinsic mitochondrial DNA dysfunction.

These results show that NSC Mito-EVs are incorporated by and restore the mitochondrial function of persistently mtDNA-depleted target cells.

### Mito-EVs are integrated in the mitochondrial network of mononuclear phagocytes via endocytosis

Mitochondrial function and immune metabolism guide the activation of mononuclear phagocytes in response to inflammatory stimuli [51]. Our recent work suggests that exogenous NSC transplants have immunomodulatory functions and inhibit the activation of pro-inflammatory mononuclear phagocytes in response to endogenous metabolic signals *in vivo* [49]. As such, to gain further insights onto the role of EVs in the immunomodulatory effects of NSCs, we next investigated whether Mito-EVs are trafficked to mononuclear phagocytes and, in so doing, affect their function.

Bone marrow-derived macrophages (Mφ) were challenged with lipopolysaccharide (LPS) to generate reactive, pro-inflammatory macrophages (Mφ^LPS^) and then treated with MitoDsRed^+^-EVs (Fig. 6a). At 6 hrs after MitoDsRed^+^-EVs treatment, 67.47% (± 0.77) of the Mφ^LPS^ showed intracellular MitoDsRed^+^ intensity via FACS analysis, which was significantly higher than resting Mφ (Fig. 6b). Co-localisation analysis with confocal high-resolution spinning disk imaging allowed us to further investigate the intracellular fate of the incorporated Mito-EVs. We found that 6.58% (± 2.00) and 10.14% (± 2.85) of Mito-DsRed^+^ Mito-EVs co-localized with either the lysosomal (LAMP1) or the peroxisomal (PMP70) markers (Fig. 6c), which suggests limited trafficking of Mito-EVs to these cellular compartments. Instead, 48.24% (± 4.43) and 17.75% (± 3.56) of Mito-DsRed^+^ mitochondria were found either attached or included within the host Mφ^LPS^ mitochondrial network (Fig. 6d and Video S1). These data show that the majority of Mito-EVs preferentially escape the lysosomal and peroxisomal pathways, while instead co-localizing with the mitochondrial network of pro-inflammatory mononuclear phagocytes.

**Fig. 6.**
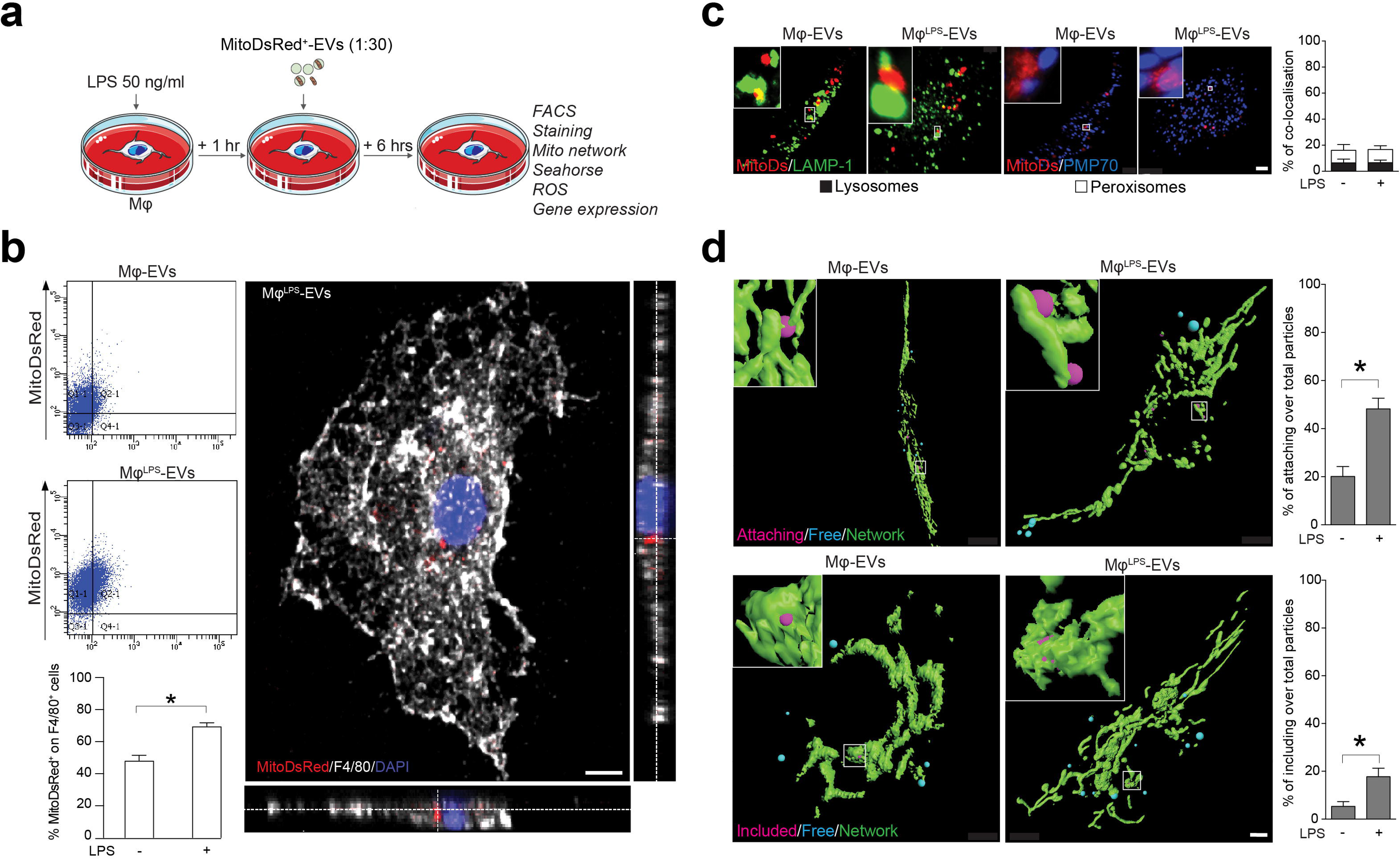
Mitochondria from NSC-derived EVs integrate in the host Mφ mitochondrial network. **a**, Experimental setup for the functional *in vitro* studies of EVs on Mφ. Mφ^LPS^ were treated with EVs spontaneously released from MitoDsRed^+^ NSCs (ratio 1:30). Uptake of MitoDSred^+^ particles and functional analyses of Mφ were assessed at 6 hrs post treatment. **b**, Flow cytometry based representative density plots of Mφ and Mφ^LPS^ at 6 hrs after treatment with MitoDsRed^+^ EVs. Data are mean % over total cells (± SEM). * p ≤ 0.05. N = 4 independent biological replicates. A representative confocal image (Orthogonal section (XY) of Z-stacks is shown) of Mφ^LPS^ treated with EVs at 6 hrs, showing uptake of MitoDSred^+^ mitochondria is also shown. Nucleus is stained with DAPI (blue). Scale bar: 3 μm. **c**, Representative spinning disk micrographs (maximum intensity projection of Z-stacks) and quantification of MitoDsRed^+^ EVs (red) co-localising with the lysosomal marker LAMP1 (green) or the peroxisomal marker PMP70 (blue) in Mφ. Data are percentage of co-localising particles over total MitoDSred+ particles (± SEM). *p < 0.05. N ≥ 10 cells per condition (2 independent experiments). Scale bars: 2 μm. **d**, Representative 3D surface (Imaris software) spinning disk images and relative quantification showing MitoDSred^+^ EVs free in Mφ (blue) or attaching (upper panels, magenta) and included (lower panels, magenta) in the mitochondrial network of Mφ (previously stained with MitoTracker Green FM). Data are percentage of either attaching or including particles over total MitoDSred^+^ particles in Mφ (± SEM). *p < 0.05. N ≥ 10 cells per condition (n = 3 independent experiments). Scale bars: 5 μm.

To further investigate the mechanism of Mito-EV incorporation into the host mitochondrial network of Mφ, we next analysed MitoDsRed^+^-EV incorporation in target Mφ^LPS^ pre-treated with either the actin-mediated phagocytosis/endocytosis inhibitor cytochalasin D (Cyto) [33, 36] or the dynamin and clathrin-mediated endocytosis inhibitors Dynasore and Pitstop 2 (D/P) [12, 41] (Fig. 7a). We found that Mφ^LPS^ showed a significantly enhanced incorporation of MitoDsRed^+^-EVs as early as 15 min after exposure (Fig. 7b,c). This effect was almost completely inhibited by the pre-treatment of Mφ^LPS^ with (D/P), suggesting that Mito-EVs are predominantly incorporated via endocytosis to be later trafficked to the host mitochondrial network.

**Fig. 7.**
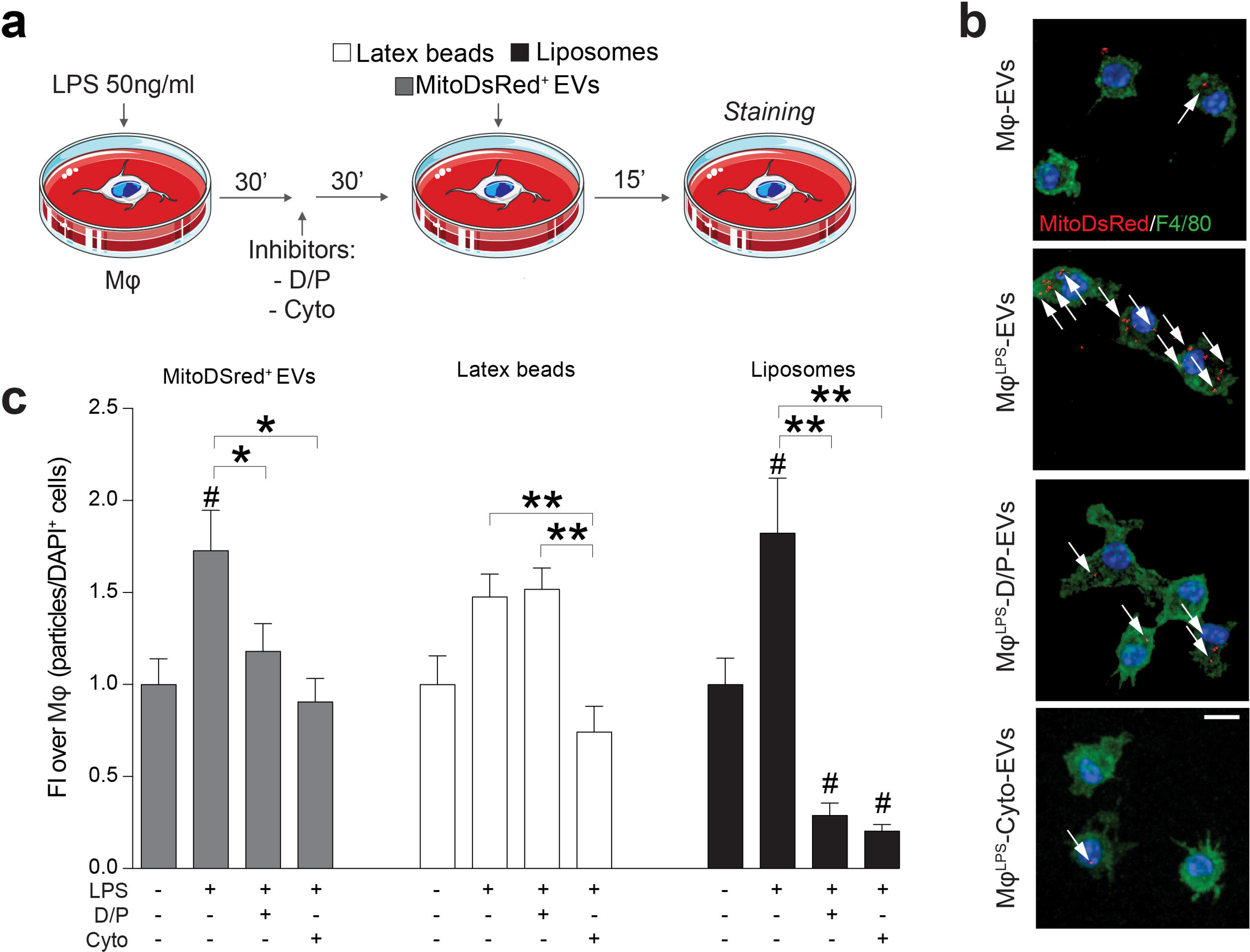
NSCs mitochondria are transferred as EVs to Mφ via endocytosis. **a**, *In vitro* experimental setup of EV uptake studies in LPS stimulated bone marrow derived Macrophages (Mφ^LPS^). Mφ^LPS^ were treated with either Cytochalsin (Cyto) or Dynasore plus Pitstop 2 (D/P) and then exposed to MitoDsRed^+^ EVs. Latex beads and liposomes were used as positive controls of phagocytosis and endocytosis, respectively. **b,c** Representative confocal microscopy images (maximum intensity projection of Z-stacks) and quantification of MitoDsRed^+^ EV (red) uptake in Mφ^LPS^ (stained for F4/80, green) in the presence or absence of endocytosis (D/P) and actin mediated phagocytosis/endocytosis (Cyto) inhibitors. Nuclei are stained with DAPI (blue). Data are particles/DAPI^+^ cells ratio and expressed as fold induction (FI) over unstimulated Mφ (± SEM). #p < 0.05 *vs* unstimulated Mφ. *p < 0.05, **p < 0.01. N ≥ 10 cells per ROI, n ≥ 8 ROI per condition (n = 3 independent experiments). Scale bars: 10 μm.

### Mito-EVs change the mitochondrial dynamics, gene expression and metabolic profile of pro-inflammatory mononuclear phagocytes

During inflammation, Mφ undergo major changes in their function and metabolism, which are associated with modifications of their mitochondrial network dynamics [34, 67]. As such, we next investigated the structure of the mitochondrial network of pro-inflammatory Mφ after EV treatment. We found that, while the stimulation with LPS promoted mitochondrial fission, the uptake of Mito-EVs and their integration into the host mitochondrial network led to a significant increase in fused mitochondria, as early as at 6 hrs after treatment (Fig. 8a).

**Fig. 8.**
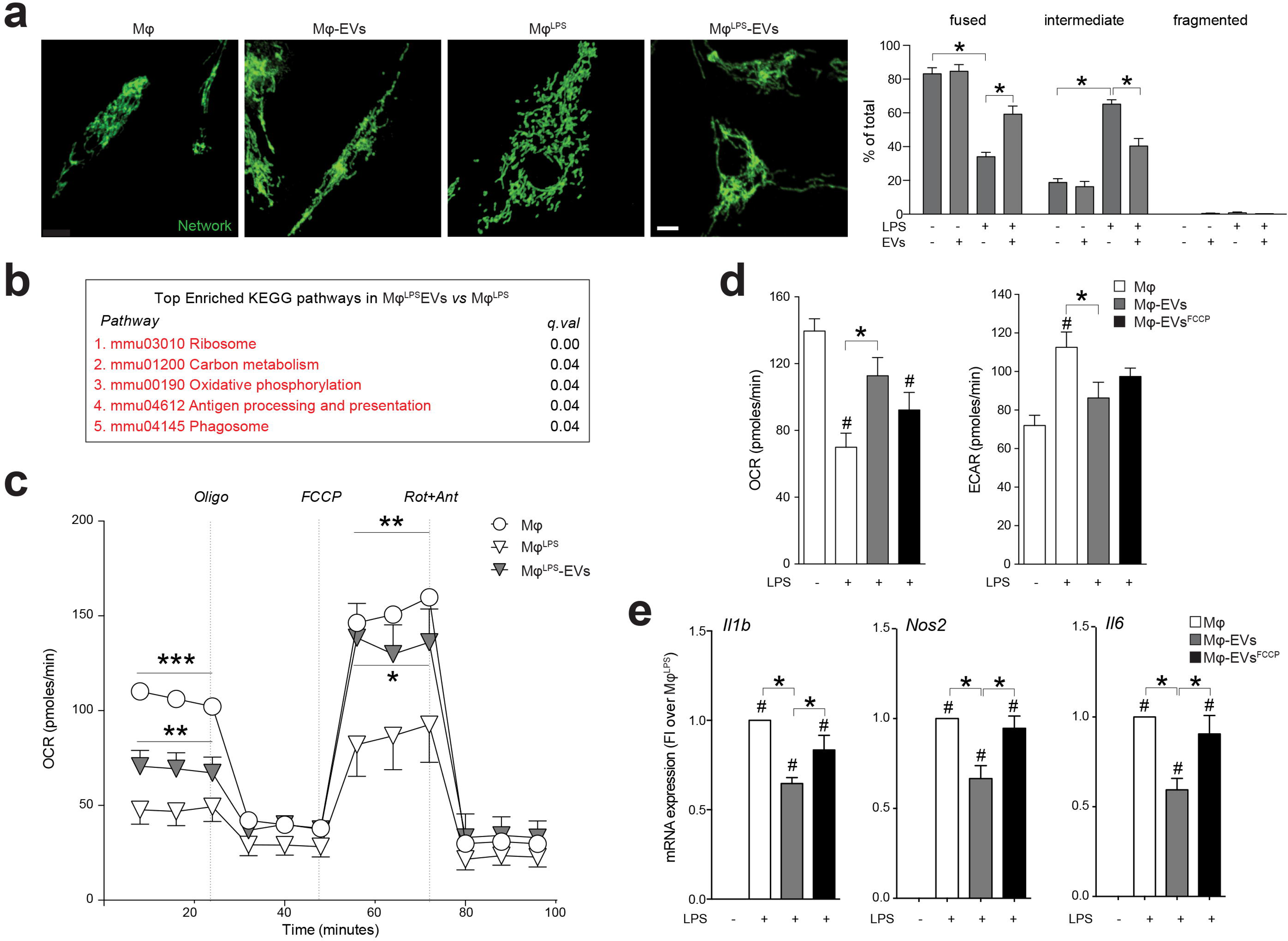
NSC-derived EVs treatment rewires Mφ^LPS^ metabolism and reduce their pro-inflammatory activation. **a**, Representative spinning disk images and quantification showing Mφ mitochondrial network labelled with TOMM20 (green) polymorphic dynamics after LPS stimulation and/or EV treatment. Data are expressed as % of Mφ displaying a fused, intermediated or fragmented network over the total number of Mφ analysed (± SEM). *p< 0.05 (n= 2 independent experiments). Scale bars: 4 μm. **b**, Top enriched KEGG pathways in genes upregulated in EV-treated *vs* untreated Mφ^LPS^ at 6 hrs. Expression data obtained by microarray analysis. **c**, XF assay of the Oxygen Consumption Rate (OCR) during a mitochondrial stress protocol of Mφ^LPS^ at 6 hrs from EVs treatment, *vs* Mφ^LPS^. Unstimulated Mφ were used as controls. Data are normalized on total protein content and expressed as mean values (± SEM). *p < 0.05, **p < 0.01, p < 0.001. N = 3 independent experiments. **d**, XF assay of the basal OCR and Extracellular Acidification Rate (ECAR) of Mφ^LPS^ at 6 hrs from treatment with EVs or treatment with EVs pre-exposed to the uncoupling agent FCCP, *vs* Mφ^LPS^. Unstimulated Mφ were used as controls. Data were normalized on total protein content and are expressed as mean values (± SEM). *p< 0.05. N = 3 independent experiments. **e**, Expression levels (qRT-PCR) of pro-inflammatory genes (*IL-1ß*, *iNOS* and *IL6*) in Mφ^LPS^ at 6 hrs from treatment with EVs or treatment with EVs pre-exposed to the uncoupling agent FCCP. Data are normalised on the housekeeping gene ß-actin and expressed mean fold induction (FI) over unstimulated Mφ (± SEM). #p< 0.05 *vs* unstimulated Mφ. *p < 0.05, **p < 0.01, ***p < 0.001. N = 2 independent experiments.

To understand the relevance of these structural changes and their functional consequences in a broader context, we analysed the gene expression profiles of Mφ^LPS^ treated with EVs using RNA expression microarrays (Table S2). Generally Applicable Gene-set Enrichment (GAGE) analysis [39] allowed us to identify specific Kyoto Encyclopedia of Genes and Genomes (KEGG) pathways upregulated in Mφ^LPS^ treated with Mito-EVs. Pathways related to ribosomes (mmu03010, q-value <0.01), carbon metabolism (mmu01200, q-value = 0.04), oxidative phosphorylation-OXPHOS (mmu00190, q-value = 0.04), antigen processing and presentation (mmu04612, q-value = 0.04) and phagosomes (mmu04145, q-value = 0.04) were all upregulated in Mφ^LPS^ treated with EVs (Fig. 8b). Interestingly, among the genes differentially expressed in the OXPHOS pathway, several genes encoding for the different subunits of the ETC were upregulated in Mφ^LPS^ after EV treatment, as shown by the Pathview diagram [38] (Fig. S2), suggesting a putative increase of cellular respiration.

As such, we next measured the oxygen consumption rate (OCR) of Mφ^LPS^ after EVs treatment, and found that the basal OCR, as well as the maximum respiratory capacity was increased in Mφ^LPS^ after EV treatment (Fig. 8c). These findings are in line with an increase in maximal respiration rate as an index of metabolic activity associated with fused mitochondrial networks [58] and they show that Mito-EVs can revert the transient mitochondrial dysfunction associated with the pro-inflammatory state of Mφ.

To prove that functional mitochondria trafficked within EVs were indeed responsible for the abovementioned changes, we treated Mφ^LPS^ with EVs that had been pre-exposed to the mitochondrial un-coupler carbonyl cyanide-4-(trifluoromethoxy)phenylhydrazone (FCCP), EV^FCCP^. While treatment with control EVs rescued the changes in the OCR and extracellular acidification rate (ECAR) induced by LPS in Mφ, treatment with EV^FCCP^ failed to do so (Fig. 8d). Moreover, control EVs, but not EV^FCCP^, succeeded in downregulating the expression of the LPS-induced pro-inflammatory cytokine genes *Il1b*, *Il6 and Nos2* (Fig. 8e) in Mφ^LPS^.

Overall these data show that NSC Mito-EVs integrate into the transiently dysfunctional host mitochondrial network of pro-inflammatory Mφ, where they re-establish physiological mitochondrial dynamics, cellular metabolism and inflammatory gene profiles.

### Transplanted NSCs transfer mitochondria to host cells *in vivo* during EAE

Previous evidence has suggested that mitochondrial transfer occurs *in vivo* and may be involved in diverse pathophysiological situations, including tissue injury and cancer progression [63]. Therefore, in order to determine if our *in vitro* findings had any functional relevance *in vivo,* NSCs or EVs were injected intracerebroventricularly (ICV) at the peak of disease (PD) into mice with MOG_35-55_-induced chronic experimental autoimmune encephalomyelitis (EAE), an animal model of multiple sclerosis.

To reliably detect NSCs and EVs *in vivo*, NSCs were previously labelled *in vitro* with both the Mito-DsRed fluorescent reporter and farnesylated (f)GFP to generate fGFP^+^/MitoDsRed^+^ NSCs. EVs spontaneously released by fGFP^+^/MitoDsRed^+^ NSCs *in vitro* were then collected and used for *in vivo* studies (Fig. 9a).

**Fig. 9.**
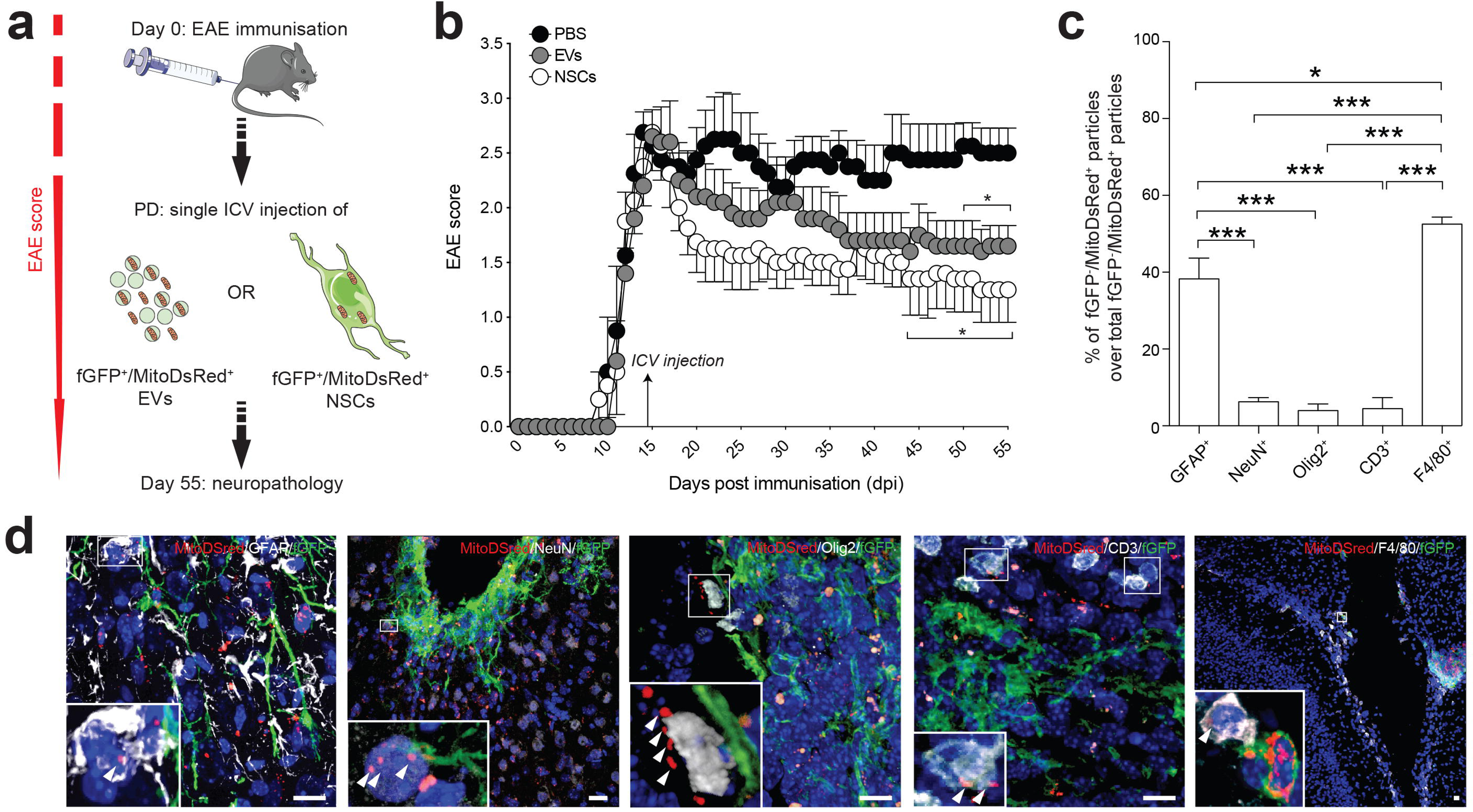
Transplanted NSCs transfer mitochondria to mononuclear phagocytes and astrocytes *in vivo* during EAE. **a**, *In vivo* experimental setup of EV and NSC treatment in EAE mice. At peak of disease (PD), mice received a single intracerebroventricular (ICV) injection of either fGFP^+^/MitoDsRed^+^ NSCs or EVs derived from fGFP^+^/ MitoDsRed^+^ NSCs (fGFP^+^/MitoDsRed^+^ EVs). Behavioural analysis was carried out daily until the end of the experiment. Neuropathology was performed at 55 days post immunisation (dpi). **b**, Behavioural outcome showing significant amelioration of the EAE score in mice treated with fGFP^+^/ MitoDsRed^+^ EVs (n = 5) and fGFP^+^/ MitoDsRed^+^ NSCs (n = 5) *vs* PBS (n = 4). *p < 0.05 *vs* PBS. **c**, Percentage of fGFP^-^/MitoDsRed^+^ particles (over total fGFP^-^/MitoDsRed^+^ particles) found in GFAP^+^ astrocytes, NeuN^+^ neurons, Olig2^+^ oligodendrocytes, CD3^+^ T cells or F4/80^+^ mononuclear phagocytes of mice treated with fGFP^+^/MitoDsRed^+^ NSCs. The majority of mitochondrial transfer events happened between transplanted NSCs and mononuclear phagocytes or astrocytes. *p < 0.05, ***p < 0.001. **d**, Representative pictures of mitochondrial transfer events detected with confocal imaging. Transfer of MitoDsRed^+^ particles (arrowheads) is shown between fGFP^+^/MitoDsRed^+^ NSCs and GFAP^+^ astrocytes, NeuN^+^ neurons, Olig2^+^ oligodendrocytes, CD3^+^ T cells or F4/80^+^ mononuclear phagocytes. Nuclei are stained with DAPI (blue). Scale bars: 20 μm.

In line with published results, a single ICV injection of NSCs resulted in a significant amelioration of EAE disease severity (from 44 days post immunisation, dpi, onwards) when compared to PBS-injected EAE mice [54]. Likewise, we found that a single ICV injection of EVs was able to significantly ameliorate EAE disability in mice (starting from 50 dpi) compared to PBS-injected EAE mice (Fig. 9b).

Following the end of the clinical observation period (55 dpi), spinal cord tissue was analysed to locate exogenous MitoDsRed^+^ immunoreactivity within the host CNS tissue. In line with the expected turnover of the MitoDsRed protein [61] we could not identify any MitoDsRed^+^ immunoreactivity in EAE mice injected with EVs. Rather, fGFP^-^/MitoDsRed^+^ mitochondria were found in EAE mice transplanted with NSCs, close to cellular grafts, suggesting the local release and transfer of mitochondria from NSCs to host CNS cells.

To determine the target cell(s) of these mitochondrial transfer events, we next quantified the number of fGFP^-^/MitoDsRed^+^ mitochondria within the three major CNS cell types (astrocytes, neurons and oligodendrocytes) and the immune cells that comprise most of the EAE inflammatory lesions (T cells and mononuclear phagocytes) (Fig. 9c). Our analysis revealed that the majority of fGFP^-^/MitoDsRed^+^ mitochondria were incorporated into mononuclear phagocytes F4/80^+^ (∼50%) and, to a lesser extent, GFAP^+^ astrocytes (∼35%). A minor fraction of fGFP^-^/MitoDsRed^+^ mitochondria (∼15%) was distributed between NeuN^+^ neurons, Olig2^+^ oligodendrocytes, or CD3^+^ T cells (Fig. 9d).

These results show that mitochondrial transfer from NSCs happens *in vivo* and that in conditions of neuroinflammation it is predominantly directed towards mononuclear phagocytes and host astrocytes.

## Discussion

Extracellular release of mitochondria and horizontal mitochondria transfer between cells are reported in several cells and organs, including the CNS [20, 53]. However, the relevance and the biological function of these phenomena are still a matter of debate.

On one hand, cells can release dysfunctional mitochondria for recycling and disposal [10, 52]. *Mitoptosis* - the selective elimination of malfunctioning mitochondria - is described in cells under conditions of severe mitochondrial stress where the occlusion of mitochondrial clusters by a membrane (“*mitoptotic body*”) allows its protrusion from the cell [40]. Similarly, *transmitophagy* - a process of transcellular degradation of damaged mitochondria through horizontal transfer - is observed between neurons and astrocytes [10] and between mesenchymal stem cells (MSCs) and macrophages [52]. In addition, it has been shown that during oxidative stress, mitochondria produce their own mitochondria-derived vesicles (MDVs) that are trafficked intracellularly either to peroxisomes [46] or to multivesicular bodies and the exosomal pathway [59]. Overall, the effect of these mechanisms is to enhance the survival of the donor cell via disposal of dysfunctional mitochondria, by routes that include unloading into neighbouring cells.

On the other hand, cells can also release intact mitochondria which retain functional properties [19, 20, 53, 56, 57, 60]. The transfer of healthy mitochondria has been demonstrated in different tissues and organs, where its main role is to maintain local homeostasis. In the CNS, astrocytes provide healthy mitochondria to damaged neurons to restore normal OXPHOS both *in vitro* and *in vivo* [20, 53]. Similarly, endothelial progenitor cells support brain endothelial energetics and barrier integrity through extracellular mitochondrial transfer [19]. MSCs exchange mitochondria to foster cytoprotection in a variety of target cells (including cardiomyocytes, endothelial cells, and corneal epithelial cells) *in vitro* [56, 57, 60] and *in vivo* [56].

Herein, we first investigated the protein content of EVs that are spontaneously released by NSCs. We found that EVs were enriched in mitochondrial proteins of the outer membrane, matrix, inner membrane and ETC, while morphological analysis confirmed the presence of entire mitochondria, as previously shown in other cellular systems [4, 6, 7]. Most importantly, we demonstrate that mitochondria spontaneously released by NSCs within EVs (Mito-EVs) have intact complexes, active complex activity, and conserved mitochondrial membrane potential and respiration. Altogether these data show that NSCs release mitochondria into the extracellular space, wherein they still harbour functional properties that can be transferred to target cells.

Unravelling the physiological significance of the mitochondrial transfer via EVs required the development of novel fluorescence and genetic mitochondrial tracking tools. To this aim, we generated NSCs lines stably expressing the MitoDsRed protein, which allowed for the expression of the fluorescent reporter in Mito-EVs. Thanks to this approach, we first provided evidence of functional mitochondria transfer from NSCs to L929 Rho^0^ cells, where Mito-EVs succeeded in reverting their intrinsic mitochondrial dysfunction and auxothropy.

Mitochondria transfer is emerging as a novel mechanism regulating the activity of the immune system. Besides the well described release of mitochondria from immune cells such as monocytes [4, 55], other immune regulatory cells modulate the activity of inflammatory cells via horizontal mitochondrial transfer [23]. As such, we next questioned whether Mito-EVs could also exert any regulatory functions on mononuclear phagocytes (Mφ) that display a transient dysfunction of mitochondria secondary to LPS stimulation.

Key to these investigations was to reveal how extracellular mitochondria enter into Mφ, which is an important step for the future development of treatment strategies designed to transfer healthy mitochondria from stem cells into immune cells. Our experiments showed that Mφ treated with EVs incorporated exogenous mitochondria preferentially via clathrin or dynamin mediated endocytosis. This is in line with previous reports showing that mitochondrial transfer events from MSCs to target cells are significantly reduced by endocytosis inhibition [57]. Most importantly, our data suggest that the internalisation of Mito-EVs is not primarily mediated by phagocytosis, but rather endocytic processes that may include micropinocytosis [35]. Indeed, we found that only a minority of the internalised Mito-EVs co-localised with either lysosomes or peroxisomes, while transferred mitochondria preferentially escaped the lysosomal pathway and integrated with the mitochondrial network of host Mφ [23].

Integration of Mito-EVs in target pro-inflammatory Mφ induced major changes in mitochondrial dynamics, gene expression profiles and metabolism. EV treatment not only increased the number of fused and intermediate mitochondria, but also induced an increase of genes related to OXPHOS, which was linked with an increase in both basal and maximal respiratory capacity of Mφ^LPS^. This is in line with previous evidence showing that inhibition of mitochondrial fission can reduce the glycolytic reprogramming of pro-inflammatory Mφ [5] and that OXPHOS-dependent ATP production can be restored by transfer of exogenous mitochondria [1, 28, 37].

It is interesting to note that the immunomodulatory effect of transferred mitochondria to immune cells reflects the activation status of their parental cells. While mitochondria derived from apoptotic cells are potent activators of innate immune responses, mitochondria derived from healthy cells are significantly less inflammatory [71]. Similarly, mitochondria from stressed monocytes, but not mitochondria from resting cells, induce type I interferon signalling in endothelial cells [55]. Here we show that NSCs release both free and membrane-embedded mitochondria with the capacity to restore OXPHOS and to reduce the pro-inflammatory gene profile of Mφ^LPS^. These effects are determined by the mitochondrial activity in parental cells rather than by the mere presence of mitochondrial content released, as shown by our experiments in which EVs treated with the mitochondrial uncoupler FCCP failed to change the gene expression profile and metabolism of recipient Mφ^LPS^. Our findings are in line with data suggesting that disrupting electron transport (or ATP synthesis) in mitochondria significantly attenuates their protective transfer effect, implying that intact OXPHOS is indispensable for this function [24].

Since our observations were based solely on *in vitro* models, we decided to investigate the relevance of mitochondrial transfer from NSCs *in vivo* in a mouse model of neuroinflammation. Mice with EAE were treated with EVs or NSCs previously labelled with fluorescent proteins to identify exogenous mitochondria and cellular grafts within the host CNS. While Mito-EVs could not be reliably identified at the time point chosen for neuropathological analysis, EAE mice treated with NSCs showed exogenous mitochondria being exchanged between the graft and host cells, suggesting a continuous production and release of mitochondria by NSCs *in vivo*. Of note, the majority of the mitochondrial transfer events was observed between NSCs and mononuclear phagocytes, strengthening the relevance of our *in vitro* findings and suggesting that similar immunoregulatory effects may be relevant also *in vivo*. Finally, a single ICV injection of EVs induced a significant amelioration of clinical deficits in EAE mice suggesting that acellular approaches, similar to NSCs grafts, may be beneficial for the treatment of neuroinflammatory disorders.

Overall, our work provides convincing proof-of-evidence that functional mitochondrial horizontal transfer occurs NSCs, paving the way for future investigations aiming at discovering the intracellular signals that link upstream mitochondrial release to downstream regulation of target cell phenotype and function. Moreover, we also support the hypothesis that NSCs mitochondrial horizontal transfer is indeed a mechanism of cell-to-cell signalling that can contribute to the immune-modulation of Mφ [69]. Future therapeutic implications of this research will need to consider strategies to pharmacologically enhance intercellular organelle transfer when desirable, such as in neuroinflammation [43, 50, 51]; or block its occurrence when it is deleterious [13].

Nonetheless, we are also fully aware of the main limitations of our work.

First, technical issues in the isolation of EVs and different nomenclatures of extracellular particles have to be considered. In this sense, a recent study by Jeppesen et al. suggests that much of the protein components of classical exosomes are absent when EVs are isolated and characterised using high-resolution density gradient fractionation and direct immunoaffinity capture [32]. However, several of the protocols used in this paper involved prolonged incubation and resuspension of immunoaffinity beads in LDS buffer, possibly causing an artefact of innate protease digestion of the EV protein cargo (including proteins that are not embedded in membranes or protected by glycosylation). Furthermore, a limited number of cell types were examined in the Jeppesen study. As such, the need for a reassessment of exosome composition and a better framework for the distinction of EVs and non-vesicular fractions is still needed.

Indeed, a second caveat of our study is the unknown nature of the NSC mitochondrial release mechanism. The release of mitochondria seems to be correlated with the metabolic state of the donor cell, as the same cell derived from different tissue sources has different mitochondrial donor properties, which are correlated with its respiratory state [48]. Cells with high mitochondrial respiration capacities are indeed associated with lower mitochondrial transfer. This is compatible with a model where donor cells optimally regulate mitochondrial transfer such that they transfer more mitochondria if they depend less on their function. Interestingly, NSCs are known to be highly glycolytic cells, which rely on glycolysis rather than OXPHOS for energy production [11]. As such, it is tempting to speculate that NSCs might have increased mitochondrial donor properties compared to other cell types, and that specific stimuli affecting their metabolism might further enhance this mechanism.

In conclusion, our work provides new insights to the contribution of mitochondria to the content and biological activity of EVs released by NSCs, suggesting that EV-mediated paracrine actions and mitochondrial transfer are two independent, but possibly interactive, pathways that allow their immunomodulatory effects [3, 66]. Overall, this work indicates that NSC mitochondrial transfer is a novel strategy of cell-to-cell signalling that might support recovery in CNS disorders.

## Acknowledgements

The authors wish to acknowledge G. Pluchino, G. Tannahill, B. Balzarotti, C. Willis and J. Brelstaff for their technical contributions and critical insights throughout the execution of the study; C. Cossetti and R. Schulte of the CIMR Flow Cytometry Core Facility for their advice and support in flow cytometry. The authors also thank A. Tolkovsky for providing the MitoDsRed plasmid and J. A. Enriquez for providing the L929-Rho^0^ cells used in this study.

This work was funded by the Italian Multiple Sclerosis Association (AISM, grant 2010/R/31 and grant 2014/PMS/4 to SP), the Italian Ministry of Health (GR08-7 to SP), the European Research Council (ERC) under the ERC-2010-StG grant agreement n° 260511-SEM_SEM, the Medical Research Council (CSF MR/P008801/1 to NJM), NHS Blood and Transplant (WPA15-02 to NJM), the Engineering and Physical Sciences Research Council, the Biotechnology and Biological Sciences Research Council UK Regenerative Medicine Platform Hub “Acellular Approaches for Therapeutic Delivery” (MR/K026682/1 to SP), the Evelyn Trust (RG 69865 to SP), the Bascule Charitable Trust (RG 75149 to SP), the Wellcome Trust (RRZA/057 RG79423 to LPJ and PRF 101835/Z/13/Z to PJL), the NIHR Cambridge BRC, and by a Wellcome Trust Strategic Award to CIMR. LPJ is supported by a senior research fellowship from FISM - Fondazione Italiana Sclerosi Multipla (cod. 2017/B/5) financed or co financed with the ‘5 per mille’ public funding. RR was funded by the german Fritz Thyssen Foundation with a one-year fellowship (40.16.0.026MN). EFV, CB and MZ are funded by a core grant of the MRC to the Mitochondrial Biology Unit (MC_UU_00015/5), ERC Advanced Grant (FP7-322424 to MZ) and NRJ-Institut de France (to MZ).

## Author contributions

Conceptualization: LPJ, JDB, NI, CF, SP; Methodology: LPJ, JDB, GM, RR, EFV, CB, AK, AVDB, TL, JW, NV, AB, CB, NI, AB, NM; Formal Analysis: LPJ, GM, RR, AK, TL, CB, AB, NM; Investigation: LPJ, GM, RR, SP; Resources: LPJ, SP; Data curation: LPJ, GM, RR, TL, CB, AK, JW, CB ; Writing - Original Draft: LPJ, GM, JAS, IB, CF, SP; Writing – Review & Editing: LPJ, GM, IB, JAS, CF, SP; Supervision: EIB, NF, CF, NM, CV, SP; Project Administration: LPJ, CV, SP; Funding Acquisition: LPJ, MZ, SP.

## Supplemental Data and Figure legends

**Fig. S1 Size distribution and proteomic analysis of EVs released by NSCs.**

**a**, Representative particle size distribution analysis of EVs by nanoparticle tracking analysis (NTA), showing bimodal distribution of particle size in the exosome size range (30-150 nm) and in the sub-micron region (20-1000 nm). Data are expressed as size distribution (nm) of EVs by particle number (% of total particle per ml) (± SD).

**b**, Comparison of EVs particle diameter (nm) and particle concentration (particle/ml) measurements using tunable resistive pulse sensing (TRPS) qNANO and NTA analysis. Data are expressed as mean values of n = 3 different experiments.

**Fig. S2 Oxidative phosphorylation genes modulated by EV in Mφ^LPS^.**

Pathview diagram showing the significantly enriched Oxidative Phosphorylation KEGG pathway. The colour scale represents the log_2_ fold change of each gene in EV-treated *vs* untreated Mφ^LPS^ as measured by microarray analysis.

**Video S1 Representative video of Mito-EVs integrating in the host Mφ^LPS^ mitochondrial network *in vitro*.**

Mito-EVs obtained from NSCs (red, arrow) is shown to interact with the host mitochondrial network of Mφ^LPS^ (stained with MitoTracker Green FM). The x/y trace and 3D rendering show a Mito-EVs (red) fusing with host mitochondria (green).

**Table S1 Data from the multiplex TMT-based proteomic experiments.**

Complete dataset (unfiltered) from multiplex TMT-based proteomic experiment illustrated in Fig 1a. Normalised, unscaled protein abundances (biological replicates 1-3), log_2_(ratios), q values and the numbers of unique peptides used for protein quantitation are shown. NSC, NSC whole cell lysates; EV, whole EV fraction; Exo, sucrose-gradient purified exosomes.

**Table S2 Data from the RNA Microarray experiments.**

Table showing the differential expression results from microarray analysis of Mφ^LPS^ treated with EVs, *vs* untreated Mφ^LPS^ at 6 hrs.

