## Supplementary material for "Neural stem cells traffic functional mitochondria via extracellular vesicles to correct mitochondrial dysfunction in target cells": Figure S1

**a**

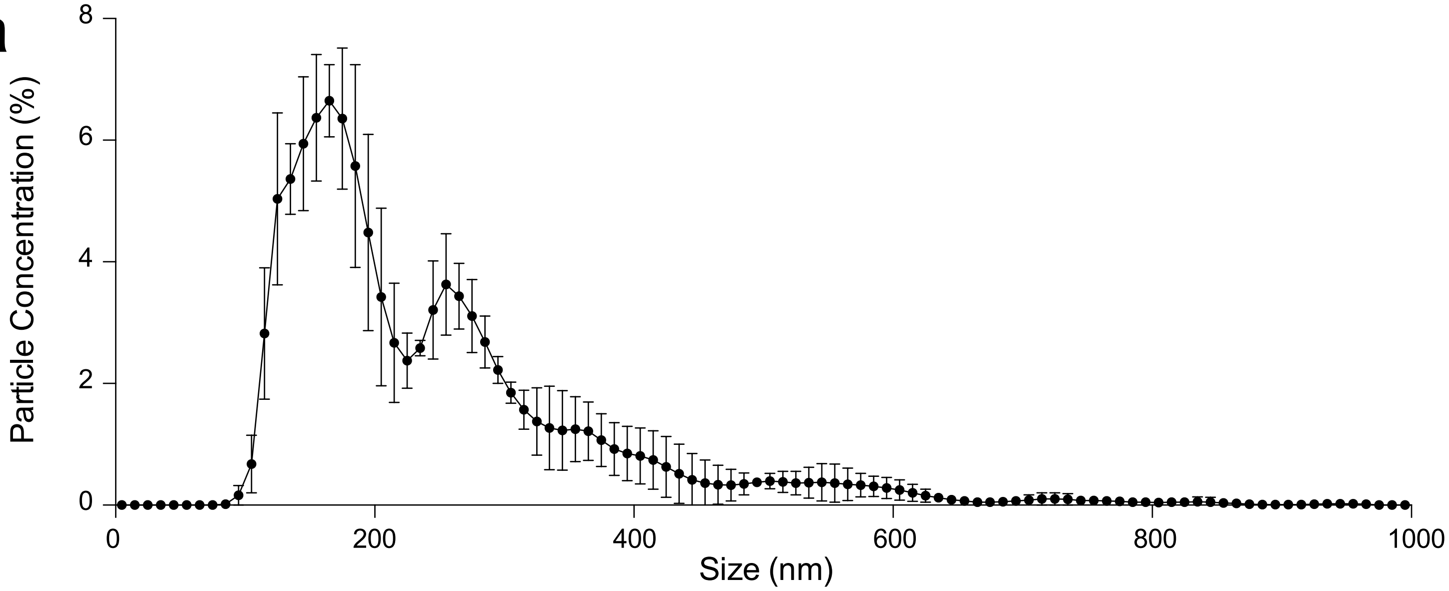


**b**

| | qNANO | NTA |
|---|---|---|
| **Particle diameter (nm)** | | |
| mean | 132.5 | 239.8 |
| mode | 80.0 | 150.8 |
| D10 | 75.5 | 126.6 |
| D50 | 107.0 | 196.3 |
| D90 | 230.5 | 366.2 |
| D90/D10 | 3.1 | 2.9 |
| **Concentration (particle/ml)** | | |
| measured | 6.0 E+09 | 9.0 E+08 |
| corrected | 6.0 E+12 | 8.0 E+11 |

**Peruzzotti-Jametti et al. Figure S1**
