## Supplementary figures and images for "Neural stem cells traffic functional mitochondria via extracellular vesicles to correct mitochondrial dysfunction in target cells"

### Figure S2

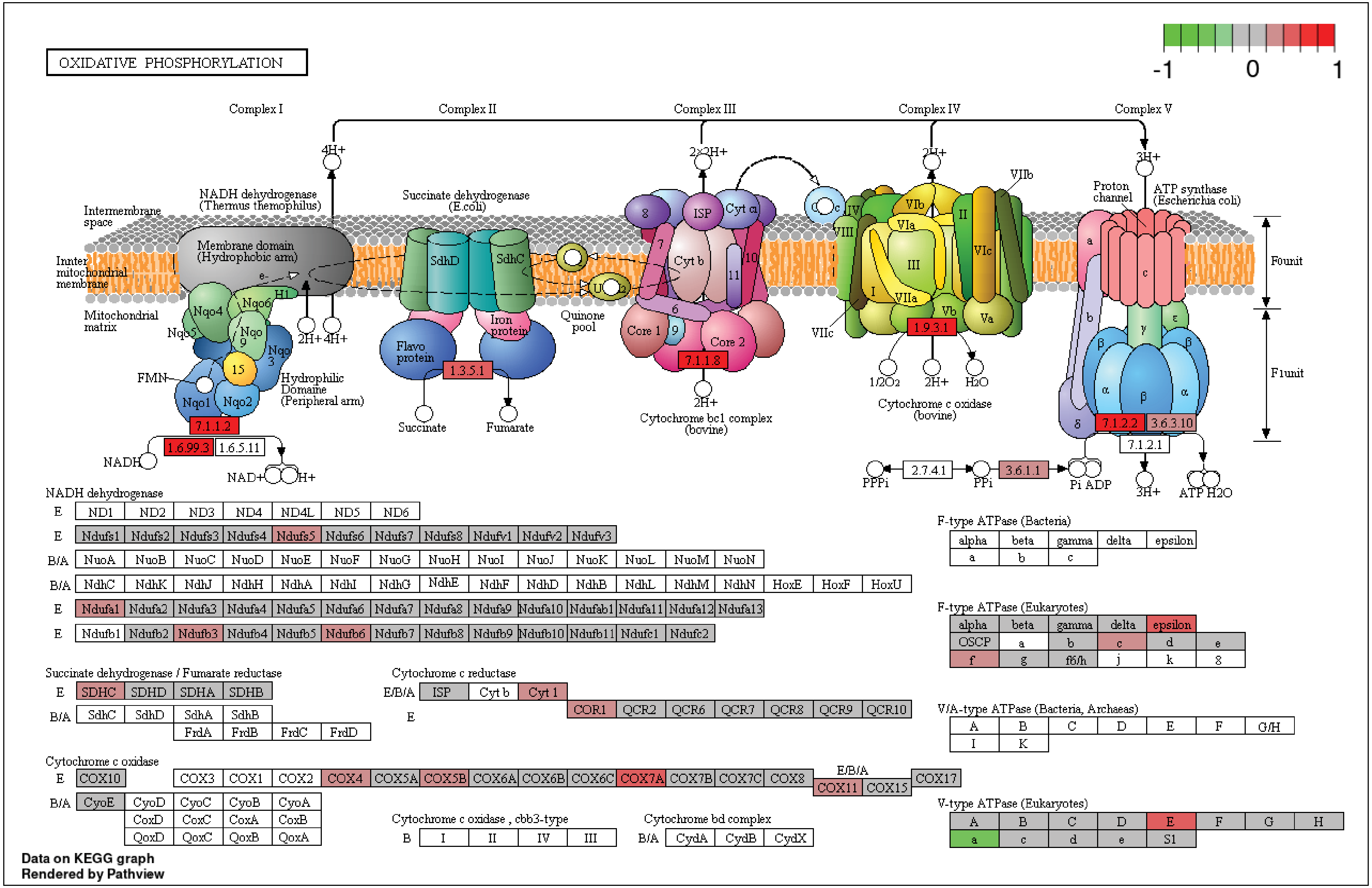


**Peruzzotti-Jametti et al. Figure S2**
